# Deciphering the antifungal mechanism of Polish ethanolic extracts of propolis against *Candida albicans*: Evidence for a multi-target mode of action

**DOI:** 10.64898/2026.04.23.720452

**Authors:** Piotr Bollin, Michał K. Pierański, Michał W. Szcześniak, Piotr Szweda

## Abstract

Propolis, a resinous product of plant origin collected by honeybees, has long been used in traditional medicine for its antimicrobial properties. However, its antifungal mechanism of action against *Candida albicans* remains incompletely understood. This study aimed to elucidate the antifungal mechanism of ethanolic extracts of Polish propolis (EEP) and a defined mixture of its key flavonoid constituents against *C. albicans*. Antifungal activity was assessed using Time-kill assay and broth microdilution method under different medium supplementation conditions. Cellular responses were analysed by fluorescence microscopy (FM) and flow cytometry (FCM) in regard to: membrane integrity, reactive oxygen species (ROS) accumulation, mitochondrial membrane potential changes, cytosolic Ca²⁺ levels, and morphological transition. Transcriptomic changes were evaluated by RNA sequencing (RNA-seq) and validated by quantitative reverse transcription PCR (RT-qPCR). EEP exhibited concentration-dependent, extract-specific antifungal activity, with selected samples showing rapid fungicidal effects. Mechanistic studies demonstrated membrane permeabilization, reactive oxygen species (ROS) accumulation, mitochondrial dysfunction, disruption of Ca²⁺ homeostasis, and inhibition of hyphal formation. Ergosterol supplementation reduced antifungal efficacy, indicating membrane sterols as primary targets. Transcriptomic analysis revealed downregulation of genes associated with DNA replication, transcription, and biosynthesis, alongside upregulation of stress-response pathways, including oxidative stress, protein folding, and mitochondrial processes. Polish propolis exerts antifungal activity through a multi-target mechanism involving membrane disruption and induction of cellular stress. Transcriptomic data indicate coordinated suppression of essential cellular functions and activation of stress-response pathways, supporting a system-level disruption of fungal homeostasis.

## 1. Introduction

*Candida albicans* is a critical fungal threat to global public health. This status was recently formalized by its inclusion in the “Critical Priority” group of the World Health Organization Fungal Priority Pathogens List (WHO, 2022). While historically recognized as the primary cause of invasive candidiasis, its threat level continues to increase. High mortality rates associated with systemic infections, ranging from 30% to over 40%, persist despite the availability of antifungal therapies (Pappas et al., 2018; Talapko et al., 2021; Denning, 2024). Clinical isolates of *C. albicans* frequently exhibit complex resistance mechanisms, including the overexpression of drug efflux pumps and alterations in drug targets, such as *ERG11* mutations, which compromise the efficacy of standard azole treatments (Bhattacharya et al., 2020). Additionally, *C. albicans* forms robust biofilms on medical devices and host tissues, increasing multidrug tolerance and serving as reservoirs for persistent and recurrent infections (Larkin et al., 2017; Lee et al., 2020B). Collectively, high mortality, evolving resistance, and biofilm-associated persistence underscore the need for novel therapeutic strategies.

Natural products, particularly those derived from traditional medicine, represent promising sources of novel antifungal compounds with unique mechanisms of action. Propolis is a well-recognized example of such a product (Kuropatnicki et al., 2013). It is a complex resinous mixture produced by honeybees (*Apis melifera*) from plant exudates, buds, and other botanical sources (Silva-Carvalho et al., 2015). Propolis extracts contain over 300 bioactive constituents, predominantly flavonoids and phenolic acids (Okińczyc et al., 2022; Ożarowski et al., 2022). However, their composition varies significantly depending on geographical origin, bee species, and seasonal factors, resulting in distinct chemical profiles and biological activities (Isidorov et al., 2016; Kasote et al., 2022). Propolis definitely belongs to the most popular (and effective) antimicrobial agents of folk medicines that was used in different regions of the world. Over the last two or three decades, there has been a renewed interest in the possibility of using this product as a substitute for antibiotics. A number of reports confirm the significant therapeutic potential of propolis, but the molecular mechanisms of its antibacterial and antifungal activity remain largely unknown (Rojczyk et al., 2020; Toreti et al., 2013).

Multiple studies suggest that ethanolic extracts of propolis (EEP) can damage the cell wall and membrane of *C. albicans*, inhibit filamentation and biofilm formation, and induce apoptosis (Corrêa et al., 2020; Cerqueira et al., 2022; Silva-Beltrán et al., 2023). However, these findings are often limited to specific extracts, strains, and experimental conditions, which hinders generalization of mechanistic conclusions and identification of specific molecular targets. Consequently, despite promising results, current data provide only a partial and sometimes inconsistent understanding of how these complex mixtures affect fungal cells and suppress virulence. Therefore, further investigation into the antifungal mechanisms of action of EEP and its individual components is warranted.

This study builds upon our previous work on the antifungal activity of ethanolic extracts of Polish propolis (Bollin et al., 2025). However, the primary objective was not to compare the composition and activity of different extracts, but to expand the understanding of antifungal mechanisms of action using previously characterized EEP, together with a mixture of components identified as key constituents of ethanolic extract of propolis (CIEEP), particularly against *C. albicans*. To achieve this, standard methods for assessing antifungal activity were employed, including evaluation of the effects of medium supplementation on EEP activity and its components, as well as time-kill assays. Intra- and extracellular changes were analyzed using fluorescence microscopy (FM) and flow cytometry (FCM). Additionally, the transcriptomic response of EEP-treated *C. albicans* cells was investigated using RNA sequencing. Changes in the relative expression of selected genes, including virulence-associated markers, were further validated by quantitative RT-PCR (RT-qPCR).

## 2. Materials and methods

### 2.1. Chemicals, strains and media

Dimethyl sulfoxide (DMSO), phosphate-buffered saline (PBS), D-(+)-glucose, Pluronic F-127, carbonyl cyanide *m*-chlorophenyl hydrazone (CCCP), oligomycin A, sodium hydroxide, potassium hydroxide, sodium citrate, 4-(2-hydroxyethyl)-1-piperazineethanesulfonic acid (HEPES), sodium chloride, potassium chloride, magnesium chloride, calcium chloride, ergosterol, bovine serum albumin (BSA), and sorbitol were obtained from Sigma-Aldrich (USA), whereas 96% ethanol and 30% hydrogen peroxide were purchased from POCH (Poland). Ultrapure water used in all experiments was obtained from a Milli-Q IQ 7005 purification system (Merck, Germany).

Ethanolic extracts of propolis - EEP 1, 8, 18, 33, 39, and 76 used in the following experiments were selected from a collection of samples analyzed in a previous study (Bollin et al., 2025) based on their antifungal activity (low or high). In addition to the EEP samples, a flavonoid mixture composed of five components, referred to as CIEEP, was included. The CIEEP mixture (MIX) consisted of pinocembrin, chrysin, galangin, pinobanksin, and pinobanksin-3-acetate (ChemFaces, China). The compounds were dissolved in 100% molecular biology grade DMSO, and the resulting solutions (10.24 µg/mL each) were combined in a 1:1:1:1:1 ratio (v/v).

The activity of EEP and the CIEEP mixture was evaluated throughout the study against the *Candida albicans* SC 5314 reference strain. *C. albicans* was routinely cultured on Sabouraud dextrose agar (Merck, Germany) or in liquid yeast extract peptone dextrose (YPD) medium (A&A Biotechnology, Poland). All antifungal susceptibility assays were performed using RPMI 1640 medium (Sigma-Aldrich, USA) adjusted to pH 7.0. The medium was supplemented with glucose to a final concentration of 2% (w/v) and buffered with 3-(N-morpholino)propanesulfonic acid (MOPS). The pH was adjusted by titration with sodium hydroxide. Before use, the medium was sterilized by filtration through 0.22 µm Millex™ PES syringe filters (Merck, Germany).

### 2.2. Time-kill assay

The antifungal activity of EEP and the CIEEP mixture was evaluated using a time-kill assay, as previously described (Bollin et al., 2025), with minor modifications. In these experiments, the initial optical density of the *C. albicans* cell suspension was adjusted to a 3 McFarland standard (approximately 5 × 10⁶ CFU/mL). Samples were collected at 0, 1.5, 3, 6, 9, 12, and 24 h, serially diluted, and plated onto Sabouraud agar. Optical density was determined using a Densilameter II (Erba Lachema, Czech Republic).

### 2.3. Study of the impact of medium supplementation on EEP/CIEEP mixture effectiveness

To determine the impact of different medium supplementation on the effectiveness of EEP and the CIEEP mixture in inhibiting growth and exerting fungicidal activity against *C. albicans*, three additives were used: 0.8 M sorbitol, 100 µg/mL ergosterol, and 50 mg/mL BSA. Standard RPMI 1640 medium supplemented with 2% glucose served as the control. Minimum inhibitory concentration (MIC) and minimum fungicidal concentration (MFC) values were determined using a two-fold serial microdilution method according to the Clinical and Laboratory Standards Institute (CLSI) guidelines (M27-A2, 2002). All procedures were performed as previously described (Bollin et al., 2025).

### 2.4. Fluorescence microscopy

For this part of the study, three fluorescent dyes were used: disodium 2,2’-ethene-1,2-diylbis[5-({4-anilino-6-[bis(2-hydroxyethyl)amino]-1,3,5-triazin-2-yl}amino)benzenesulfonate] (calcofluor white, CFW), propidium iodide (PI), and MitoTracker™ Green (MTG), all obtained from Thermo Fisher Scientific (USA). An overnight culture of *C. albicans* in YPD medium was diluted fivefold with microfiltered 0.85% NaCl solution and centrifuged for 5 min at 10,000 × g. After removal of the supernatant, the cells were resuspended in RPMI 1640 medium supplemented with 192 µg/mL of EEP 76 or the CIEEP mixture. Medium containing ethanol (for EEP) or DMSO (for the CIEEP mixture) at concentrations corresponding to those in the tested samples was used as a control.

Cell suspensions were incubated with EEP or the CIEEP mixture for 3 h at 37 °C with shaking at 180 rpm. After incubation, the samples were centrifuged and resuspended in microfiltered 0.85% NaCl solution. Fluorescent dyes were added to final concentrations of 62.5 µM (CFW), 3 µM (PI), and 200 nM (MTG). All samples were incubated for 30 min in the dark at room temperature, then centrifuged and resuspended in 0.85% NaCl. The prepared suspensions were mounted on microscope slides and covered with a coverslip.

Fluorescence images were acquired using an Olympus BX60 epifluorescence microscope (Olympus Corporation, Japan) equipped with an XC50 CCD camera and plan fluorite objectives (air, 20×; oil immersion, 60× and 100×). Images were captured using CellSens Standard software (Olympus Corporation, Japan) with automatic full-region exposure, no compensation, and a gain setting of 10 dB.

### 2.5. Flow cytometry (FCM) analysis of the effect of EEP and CIEEP mixture on *C. albicans* cells

#### 2.5.1. Preparation of the cell suspension, fluorescent dyes, media, and the research equipment

For the experiments, an overnight culture of *C. albicans* in YPD medium was used. After incubation, the cells were centrifuged for 5 min at 10,000 × g, the supernatant was removed, and the resulting pellet was resuspended in PBS buffer or RPMI 1640 medium (supplemented with 2% glucose) to obtain suspensions corresponding to a 3 McFarland standard. Tested substances (EEP or the CIEEP mixture) were added to cell suspensions in RPMI medium to reach final concentrations of 128, 256, or 512 µg/mL. Control samples consisted of cell suspensions containing ethanol or DMSO at concentrations corresponding to those in the tested samples. All suspensions were incubated with shaking for 3 h at 37 °C and 180 RPM.

The citrate buffer used was a 50 mM sodium citrate solution supplemented with 2% glucose and adjusted to pH 5. Hank’s balanced salt solution (HBSS) consisted of 10 mM HEPES, 140 mM NaCl, 5 mM KCl, 10 mM glucose, 1 mM MgCl₂, and 1.8 mM CaCl₂, with pH adjusted to 7.2.

Fluorescent dyes: propidium iodide (PI), 2′,7′-dichlorodihydrofluorescein diacetate (H₂-DCFDA), rhodamine 123 (Rh123), and Fluo-3 AM were obtained from Thermo Fisher Scientific (USA). Cytometric measurements were performed using an Agilent NovoCyte 2060R flow cytometer equipped with a NovoSampler Pro NS200 (Agilent Technologies Inc., USA). Samples were analyzed in sterile 5 mL (12 × 75 mm) polystyrene round-bottom tubes (Corning Inc., USA).

#### 2.5.2. Cell membrane integrity

After incubation with EEP or the CIEEP mixture, samples were centrifuged for 5 min at 10,000 × g and washed with 0.85% NaCl solution. PI was added to a final concentration of 3 µM, and samples were incubated for 30 min at 37 °C with shaking at 180 rpm. Following incubation, samples were centrifuged, resuspended in 0.85% NaCl solution, and fluorescence was measured using a flow cytometer.

#### 2.5.3. Formation of reactive oxygen species (ROS)

For this analysis, cell suspensions in PBS buffer were first incubated for 30 min at 37 °C with 5 µM H₂-DCFDA. After dye loading, samples were centrifuged for 5 min at 10,000 × g, the supernatant was removed, and cells were exposed to EEP or the CIEEP mixture. The positive control consisted of cells treated with 1% hydrogen peroxide. After incubation, samples were centrifuged, resuspended in 0.85% NaCl solution, and fluorescence was measured using a flow cytometer.

#### 2.5.4. Mitochondrial membrane hyperpolarization

After incubation with EEP or the CIEEP mixture, samples were centrifuged and washed with citrate buffer. Rh123 was added to a final concentration of 25 µM, and samples were incubated for 30 min at 37 °C. After incubation, samples were centrifuged, resuspended in 0.85% NaCl solution, and fluorescence was measured using a flow cytometer. Additional controls included cells treated with CCCP (0.5 µM) and oligomycin A (0.1 µg/mL) to validate fluorescence shifts.

#### 2.5.5. Changes of intracellular Ca^2+^ ion concentration

After incubation with EEP or the CIEEP mixture, samples were centrifuged and washed with HBSS buffer. Fluo-3 AM, dissolved in 20% Pluronic F-127, was added to a final concentration of 5 µM, and samples were incubated for 30 min at 37 °C. After incubation, samples were centrifuged, the supernatant was removed, and cells were resuspended in 0.85% NaCl solution for flow cytometric analysis. The positive control consisted of cells exposed to alkaline stress using 3 M KOH, as described by Wang et al. (2011).

#### 2.5.6. Morphological changes of *C. albicans* cells

The effect of EEP and its components on morphological changes in *C. albicans* cells, i.e., the transition from blastospores to larger hyphal or pseudohyphal forms, was analyzed using flow cytometry based on forward scatter height (FSC-H). Samples were prepared as described above, without fluorescent dyes. An empirical FSC-H threshold (AU = 1.9) was established based on comparison of signal distributions obtained from cells grown overnight in YPD medium (non-inducing conditions) and cells incubated for 3 h in RPMI medium (hyphae-inducing conditions). This threshold was used to distinguish the blastospore population (signals below AU = 1.9) from the population enriched in hyphal/pseudohyphal forms (signals above AU = 1.9). (Figure S1 in Supplementary data).

### 2.6. RNA isolation

An overnight culture of *C. albicans* was diluted 50-fold in RPMI medium and incubated for 5 h at 37 °C with shaking at 180 rpm to reach the logarithmic growth phase. The culture was then centrifuged at 10,000 × g for 5 min and resuspended in fresh RPMI medium. Aliquots of 2 mL were incubated for 3 h with 192 µg/mL of the respective EEP or CIEEP mixture. Control samples were treated with equivalent volumes of either 70% ethanol (for EEPs) or DMSO (for the CIEEP mixture).

Following incubation, cultures were centrifuged, resuspended in 800 µL of Fenozol, and incubated overnight at 4 °C. RNA was then isolated according to the manufacturer’s protocol using the Bead-Beat Total RNA Mini Kit (A&A Biotechnology, Poland). Samples were further purified using the Cut&Go gDNA Removal Kit (EurX Sp. z o.o., Poland), following the manufacturer’s instructions. RNA concentration and purity were assessed using a BioSpec-nano spectrophotometer (Shimadzu Corp., Japan). RNA samples were stored at −80 °C or kept on ice during handling.

### 2.7. RNA sequencing

#### 2.7.1. Preparation of samples

After exposure of cells to EtOH (for control samples) or EEP 76, the obtained RNA was treated with DNase I (Qiagen, Germany) for 10 min at 25 °C. Subsequently, sodium acetate (pH 5.2) was added to a final concentration of 0.3 M, followed by the addition of 2.5 volumes of ice-cold absolute ethanol. Samples were incubated at −20 °C overnight and centrifuged for 15 min at 15000 × g at 4 °C. The RNA pellet was washed twice with cold 70% ethanol, air-dried for 5 min at 37 °C, and resuspended in RNase-free water.

Samples were then stored at -80 °C and shipped on dry ice to Novogene GmbH (Germany) for RNA sequencing. Poly(A)-enriched mRNA libraries were prepared. At least 6 Gb of raw data per sample were generated using the NovaSeq X Plus platform (paired-end 150 bp reads, PE150).

#### 2.7.2. RNA-seq data analysis

##### 2.7.2.1. Quality control and read preprocessing

The initial quality assessment of raw sequencing reads was performed using FastQC v0.12.1 (Andrews, 2010) and MultiQC v1.18 (Ewels et al., 2016). Adapter removal and quality trimming were carried out with BBDuk2, a component of the BBTools suite v37.02 (Bushnell, 2014), using the following parameters: sliding window quality trimming (qtrim = w) with a minimum quality threshold of 20 (trimq=20), minimum average read quality of 10 (maq = 10), adapter matching with k-mer size of 23 (k=23), minimum k-mer length of 11 (mink = 11), Hamming distance of 1 (hdist = 1), and a minimum read length of 100 bp (minlength = 100). Adapter sequences were referenced from the BBTools built-in adapter database. Ribosomal RNA (rRNA) reads were subsequently removed by aligning trimmed reads against *C. albicans* rRNA sequences obtained from Ensembl Fungi (Cunningham et al., 2022) using Bowtie2 v2.5.3 (Langmead & Salzberg, 2012) in fast alignment mode, retaining only unaligned (non-rRNA) read pairs for further analysis.

##### 2.7.2.2. Gene expression quantification

Cleaned and filtered reads were quantified at the gene level using RSEM v1.3.1 (Li & Dewey, 2011) with Bowtie2 as the underlying aligner. RSEM was run in paired-end mode, and expected read counts were extracted for each gene. A count matrix suitable for differential expression analysis was then constructed from the RSEM gene-level output files using a custom Python script.

##### 2.7.2.3. Differential expression analysis

Differential gene expression between the treated (SCP) and untreated control (SCE) samples was assessed using the DESeq2 v1.38.3 package (Love et al., 2014) in R v4.2.1 (R Core Team, 2023). Raw count data were normalized and tested for differential expression using the default DESeq2 workflow, with the control condition set as the reference level. Genes with a Benjamini-Hochberg adjusted p-value (padj) < 0.05 were considered significantly differentially expressed. Results were annotated with gene names and functional descriptions retrieved from Ensembl Fungi BioMart (Cunningham et al., 2022).

##### 2.7.2.4. Data visualization

Principal component analysis (PCA) was performed on regularized log-transformed (rlog) expression values using the plotPCA function from DESeq2. PCA plots were generated using ggplot2 v3.4.2 (Wickham, 2016) with ggrepel v0.9.3 (Slowikowski, 2023) for non-overlapping sample labels. Sample-to-sample distance clustering was computed using Euclidean distances on rlog-transformed data and visualized as a heatmap using the heatmap.2 function from gplots v3.1.3 (Warnes et al., 2022) with the GnBu color palette from RColorBrewer (Neuwirth, 2022). MA plots were generated using the plotMA function from DESeq2. Expression heatmaps of the top 50 most significantly differentially expressed genes (ranked by adjusted p-value) were generated using pheatmap v1.0.12 (Kolde, 2019). Expression values were rlog-transformed and subsequently converted to Z-scores (row-wise standardization). Hierarchical clustering of genes (rows) was performed, while sample columns were displayed in a fixed order. Gene names were mapped from systematic locus identifiers using annotation data obtained from Ensembl Fungi BioMart.

##### 2.7.2.5. Functional enrichment analysis

Gene Ontology (GO) enrichment analysis was performed separately for up-regulated and down-regulated genes (log2 fold change > 2 and < -2, respectively) using the enricher function from the clusterProfiler v4.6.2 package (Wu et al., 2021) with custom GO term-to-gene mappings derived from *C. albicans*-specific annotation data obtained from Ensembl Fungi BioMart (Cunningham et al., 2022). Enrichment was tested independently for three GO categories: Biological Process (BP), Molecular Function (MF), and Cellular Component (CC). The Benjamini-Hochberg method was used for multiple testing correction, with both p-value and q-value cutoffs set at 0.05. For visualization, the top 15 enriched BP terms per direction were selected and assigned to broad functional categories (e.g., rRNA processing, DNA repair/replication, Proteostasis/UPS, Stress response, Mitochondria/Energy) based on GO term descriptions, to highlight biological differences between up- and down-regulated gene sets. Results were visualized as a combined dot plot using ggplot2 v3.4.2 and cowplot v1.1.1 (Wilke, 2024).

### 2.8. Reverse transcription (RT)

To synthesize cDNA from isolated RNA, the Smart First Strand cDNA Synthesis Kit (EurX Sp. z o.o., Poland) was used with random hexamers as primers. All steps of reaction mixture preparation were performed according to the manufacturer’s protocol. Reverse transcription was carried out using a Mastercycler Gradient 5331 (Eppendorf AG, Germany) under the following conditions: 10 min at 25 °C, 50 min at 50 °C, and 5 min at 85 °C. cDNA was stored at −20 °C and kept on ice during handling.

### 2.9. qPCR

Quantitative PCR (qPCR) of cDNA samples was performed using Fast SG qPCR Master Mix (EurX, Poland) according to the manufacturer’s instructions. Reactions were carried out using a LightCycler 96 instrument (Roche, Switzerland). The amplification and detection of specific products were performed under the following conditions: an initial denaturation step at 95 °C for 10 min, then, 40 cycles of amplification consisting of 95 °C for 15 s and 60 °C for 1 min. This was followed by a melting curve analysis, performed by heating to 95 °C for 10 s, cooling to 55 °C for 1 min, and increasing the temperature to 97 °C. Primers for all analyzed genes (forward and reverse) were designed using the Primer-BLAST tool (Ye et al., 2012) and used at experimentally optimized concentrations (Table S1). Primers were synthesized by Oligo.pl (IBB PAN, Poland).

Relative gene expression was quantified using the Pfaffl method (Pfaffl, 2001), taking into account individual qPCR efficiencies determined from standard curves. Two reference genes (*ACT1* and *RIP1*) were used for normalization. These genes were selected based on expression stability ranking (Table S2) obtained using RefFinder software (Xie et al., 2023), with cells treated with EEP 76 used for stability assessment. For each biological replicate, the geometric mean of reference gene expression (E^−Cq) was calculated and used to normalize target gene expression.

Normalized expression levels of EEP- or CIEEP-treated samples were expressed relative to the corresponding ethanol (for EEP) or DMSO (for the CIEEP mixture) control for each biological replicate. For the CIEEP mixture, only *ACT1* was used for normalization due to the instability of *RIP1* expression under these conditions.

### 2.10. Data analysis

All experiments were performed in triplicate, and data are presented as mean ± SD. Graphs were prepared using GraphPad Prism® 8.0.2 (GraphPad Software Inc., USA). Where applicable, statistical analysis was performed using two-way ANOVA in the same software. Flow cytometry data was analyzed using FlowJo v10.

## 3. Results and discussion

### 3.1. Antifungal activity of EEP and CIEEP mixture over time

At the lowest concentration (128 µg/mL), EEP 33 demonstrated the highest fungicidal activity among all tested samples, achieving approximately a 3 log₁₀ reduction by 9 h and sustaining this effect through 12 h, followed by partial regrowth at 24 h. EEP 76 and the CIEEP mixture exhibited primarily fungistatic effects at this concentration, maintaining relatively stable cell counts throughout the exposure period, with reductions ranging from 0.3 to 1.5 log₁₀ at different time points. The remaining extracts (EEP 1, EEP 8, EEP 18, and EEP 39) did not show significant antifungal activity at this concentration.

At 256 µg/mL, fungicidal activity increased markedly for several samples. EEP 33 and EEP 76 reduced viable counts below the limit of detection (2 log₁₀ CFU/mL) by 6 h and maintained this effect up to 24 h. EEP 8 exhibited time-dependent killing, reaching approximately a 2 log₁₀ reduction by 3 h and achieving a 3–4 log₁₀ reduction by the end of incubation, although viable cells remained detectable. The CIEEP mixture showed rapid fungicidal activity, achieving approximately a 4 log₁₀ reduction within 1.5 h and maintaining a reduction of 3.5-4 log₁₀ throughout the remaining exposure period. EEP 39 demonstrated moderate activity, with a 1-2 log₁₀ reduction, whereas EEP 1 and EEP 18 remained largely ineffective at this concentration.

At the highest concentration (512 µg/mL), rapid and near-complete fungicidal activity was observed for multiple extracts. EEP 8, EEP 33, EEP 76, and the CIEEP mixture achieved almost complete eradication by 3 h. In contrast, EEP 1 and EEP 18 exhibited primarily fungistatic effects throughout the exposure period.

These results demonstrate concentration-dependent and extract-specific antifungal activity, with EEP 33, EEP 76, and the CIEEP mixture showing the most potent fungicidal effects against *C. albicans*.

### 3.2. Addition of exogenous ergosterol or BSA to the medium affects the EEP and CIEEP mixture effectiveness

To investigate the impact of growth medium additives on the antifungal effectiveness of EEP and the CIEEP mixture, supplementation with ergosterol, sorbitol, or BSA was evaluated. Ergosterol (ergosta-5,7,22-trien-3β-ol) is the major sterol in fungal cell membranes and plays a critical role in regulating membrane fluidity, structure, and cellular processes, making it essential for cell viability (Rodrigues, 2018). Its addition to the growth medium can indicate whether ergosterol is a molecular target of active components present in EEP, as reflected by changes in effective concentrations. In this study, ergosterol supplementation resulted in at least a two-fold increase in both MIC and MFC values for all tested samples compared to the control medium. The observed changes might be due to either direct interaction of the compounds present in the tested extracts/mixture with ergosterol and their neutralization before they affect the cells, or the result of cells compensating the molecularly induced deficiency of that crucial sterol by absorption from the environment. However, both of these hypothesis require further investigation.

In contrast, sorbitol supplementation did not result in any detectable changes in MIC or MFC values for any of the tested samples. Sorbitol acts as an osmotic stabilizer for fungal cells with compromised cell walls (Gucwa et al., 2018). When an antifungal agent primarily targets the cell wall, the presence of sorbitol protects cells from osmotic lysis, typically leading to an increase in MIC values compared to standard medium (Leite et al., 2014). The lack of such an effect in this study suggests that the fungal cell wall is unlikely to be the primary target of EEP or the CIEEP mixture.

The observations regarding ergosterol and sorbitol are consistent with previous findings for Polish propolis extracts (Gucwa et al., 2018), where exogenous ergosterol increased MIC values against *C. albicans* SC 5314 by up to sixteen-fold, indicating a strong interaction between propolis components and membrane sterols. In contrast, sorbitol did not significantly affect EEP activity, supporting the conclusion that the cell wall is not the primary target. However, Pippi et al. (2015) reported different results for a benzophenone-enriched fraction of Brazilian red propolis, where ergosterol supplementation did not affect susceptibility, while sorbitol increased MIC values up to 64-fold, suggesting a cell wall-targeting mechanism. These discrepancies may be explained by substantial differences in the chemical composition of European and Brazilian propolis (Fabris et al., 2013), potentially resulting in distinct mechanisms of action.

Supplementation of the medium with bovine serum albumin (BSA) rendered all tested samples ineffective, with MIC and MFC values exceeding the analyzed concentration range. BSA is a well-characterized globular protein commonly used as a model for protein-binding interactions due to its structural similarity to many serum and cytoplasmic proteins (Majorek et al., 2012). The addition of BSA was intended to assess whether antifungal activity is reduced due to protein binding. The observed increase in MIC and MFC values suggests that EEP components may exhibit strong protein-binding affinity, leading to reduced activity in protein-rich environments. While protein-polyphenol interactions are well documented in food chemistry (Xia & Kai, 2012; Ansari et al., 2015), their relevance to antifungal therapy remains less explored. These findings may indicate a potential limitation of EEP-based treatments under physiological conditions, where high protein concentrations could reduce bioavailability.

**Table 1.** MIC and MFC values of EEP and CIEEP mixture estimated against *C. albicans* SC 5314 using RPMI 1640 medium with different additives. Numbers with a greater-than symbol (>) indicate values outside of the analyzed concentration range. MIC/MFC values determined using medium without any additive were previously published in Bollin et al. (2025).

| Medium additive | MIC/MFC value [µg/mL] |  |  |  |  |  |  |  |  |  |  |  |  |  |
| --- | --- | --- | --- | --- | --- | --- | --- | --- | --- | --- | --- | --- | --- | --- |
|  | EEP 1 |  | EEP 8 |  | EEP 18 |  | EEP 33 |  | EEP 39 |  | EEP 76 |  | MIX |  |
|  | MIC | MFC | MIC | MFC | MIC | MFC | MIC | MFC | MIC | MFC | MIC | MFC | MIC | MFC |
| - | >256 | >256 | 256 | 256 | >256 | >256 | 128 | 128 | 256 | 256 | 256 | 256 | 64 | 128 |
| sorbitol | >256 | >256 | 256 | 256 | >256 | >256 | 128 | 128 | 256 | 256 | 256 | 256 | 64 | 128 |
| ergosterol | >256 | >256 | >256 | >256 | >256 | >256 | >256 | >256 | >256 | >256 | >256 | >256 | 256 | >256 |
| BSA | >256 | >256 | >256 | >256 | >256 | >256 | >256 | >256 | >256 | >256 | >256 | >256 | >256 | >256 |

### 3.3. FCM assessments of EEP/CIEEP mixture-treated *C. albicans*

The FCM study was based on the detection and estimation of shifts in fluorescence intensity of stained cells relative to the untreated control sample.

PI is a fluorescent molecule that binds to DNA but is unable to passively enter cells with an intact plasma membrane. Therefore, PI uptake is commonly used to distinguish dead cells with permeable plasma membranes from live cells with intact membranes, regardless of the cause of death (Crowley et al., 2016). H₂-DCFDA is a broad-spectrum intracellular ROS probe that, after esterase cleavage and oxidation to DCF, is detected as green fluorescence, with intensity proportional to intracellular oxidant levels (Ameziane-El-Hassani & Dupuy, 2013). Fluo-3 AM is a membrane-permeant acetoxymethyl ester derivative of the Ca²⁺ indicator Fluo-3. After entering cells, intracellular esterases cleave the AM groups, trapping the dye in the cytosol, where Ca²⁺ binding leads to a marked increase in fluorescence, enabling monitoring of intracellular Ca²⁺ levels (Saavedra Molina et al., 1990; Lee et al., 2020A). Rhodamine 123 (Rh123) is a cell-permeant, cationic fluorescent dye used to assess mitochondrial membrane potential (ΔΨm), as it accumulates selectively in active mitochondria due to their negative inner membrane potential (O’Connor et al., 1988).

#### 3.3.1. Membrane permeabilization

Exposure of *C. albicans* to EEP or the CIEEP mixture for 3 h resulted in a substantial increase in the percentage of PI-positive cells compared to the control, indicating compromised cell membrane integrity (Fig. 2A). The CIEEP mixture exhibited the highest activity, causing extensive membrane permeabilization with 95.5% and 91.5% PI-positive cells at 128 and 256 µg/mL, respectively. Among the extracts, EEP 33 demonstrated a clear dose-dependent relationship, with compromised cell populations increasing from 38.6% at 128 µg/mL to 79.6% at 512 µg/mL. EEP 1 and EEP 8 also showed significant membrane damage, reaching 64.1% and 53.5% PI-positive cells at the highest concentration tested, respectively. These findings are consistent with previous studies describing propolis as affecting the fungal cell envelope, leading to increased permeability, leakage of intracellular constituents, and eventual cell death (D’Auria et al., 2003; Corrêa et al., 2020). A discrepancy was observed for EEP 39 and EEP 76 at the highest concentration (512 µg/mL), where the percentage of PI-positive cells plateaued or decreased compared to 256 µg/mL (e.g., EEP 39 decreased from 60.0% to 55.1%). However, the time-kill assay results (Fig. 1) confirmed that these concentrations resulted in total eradication of the fungal population after 3 h. This suggests that the high fungicidal activity at 512 µg/mL may lead to rapid and extensive cell lysis. The resulting cellular debris or severely damaged cells may be excluded from detection or may not retain the dye efficiently, leading to underestimation of lethality when measured solely by PI fluorescence (Crowley et al., 2016). Consequently, while FCM reflects membrane damage at sub-lethal concentrations, it may underestimate total cell death under conditions of extensive cellular disintegration.

**Figure 1.**
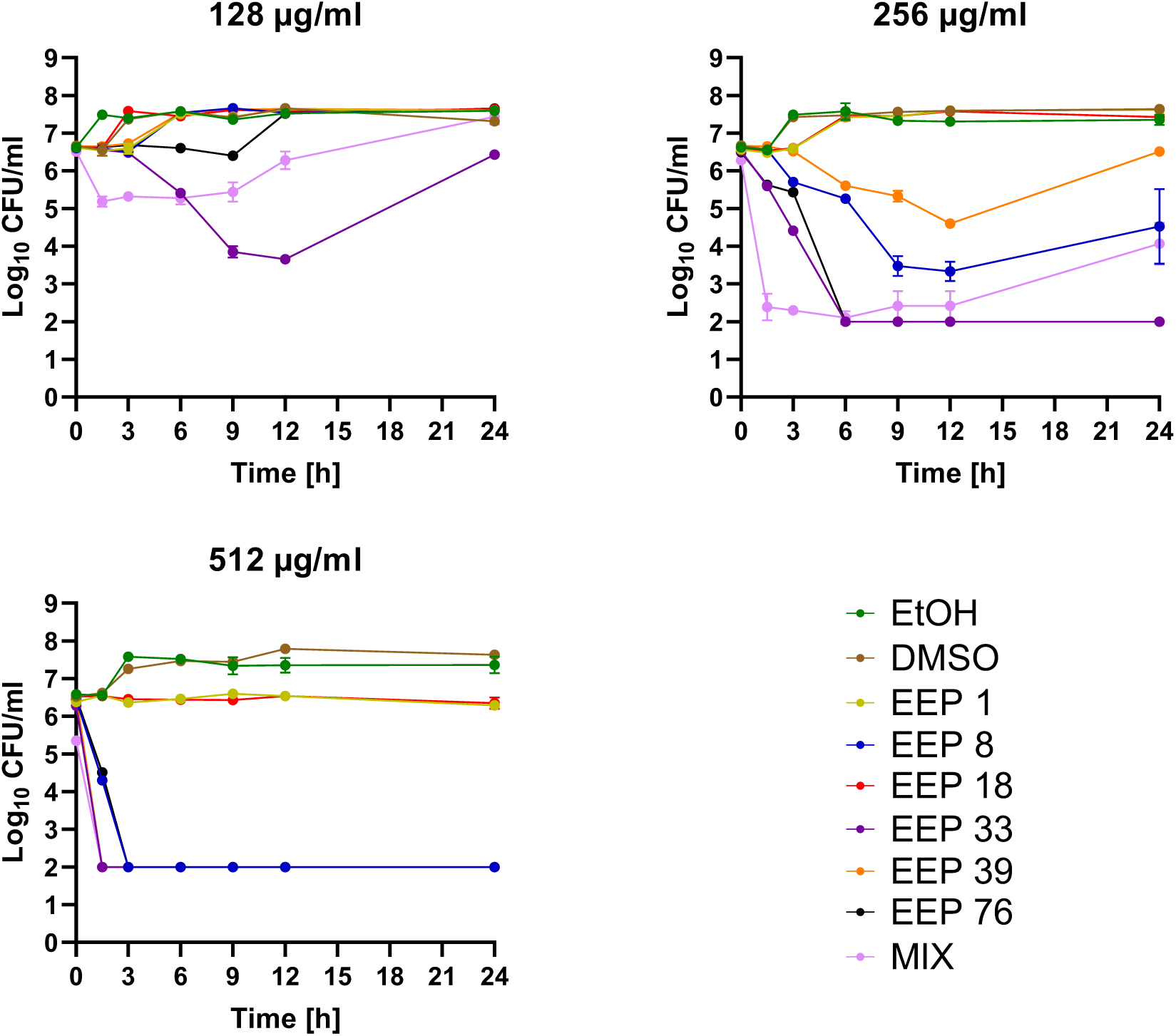
Survivability of *C. albicans* SC 5314 cells over 24 h exposed to different EEPs and CIEEP mixture in concentrations of 128, 256, and 512 µg/mL

**Figure 2.**
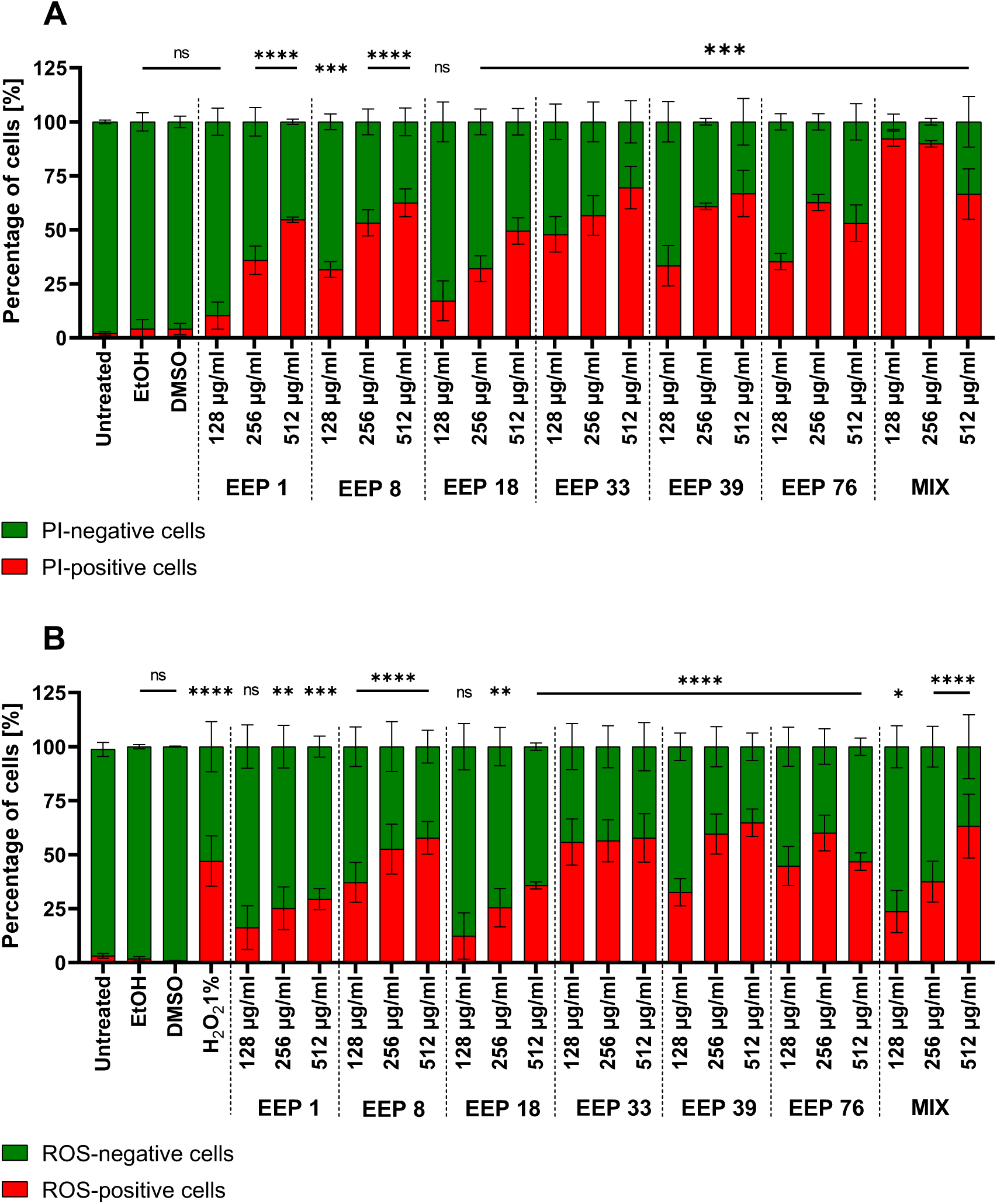
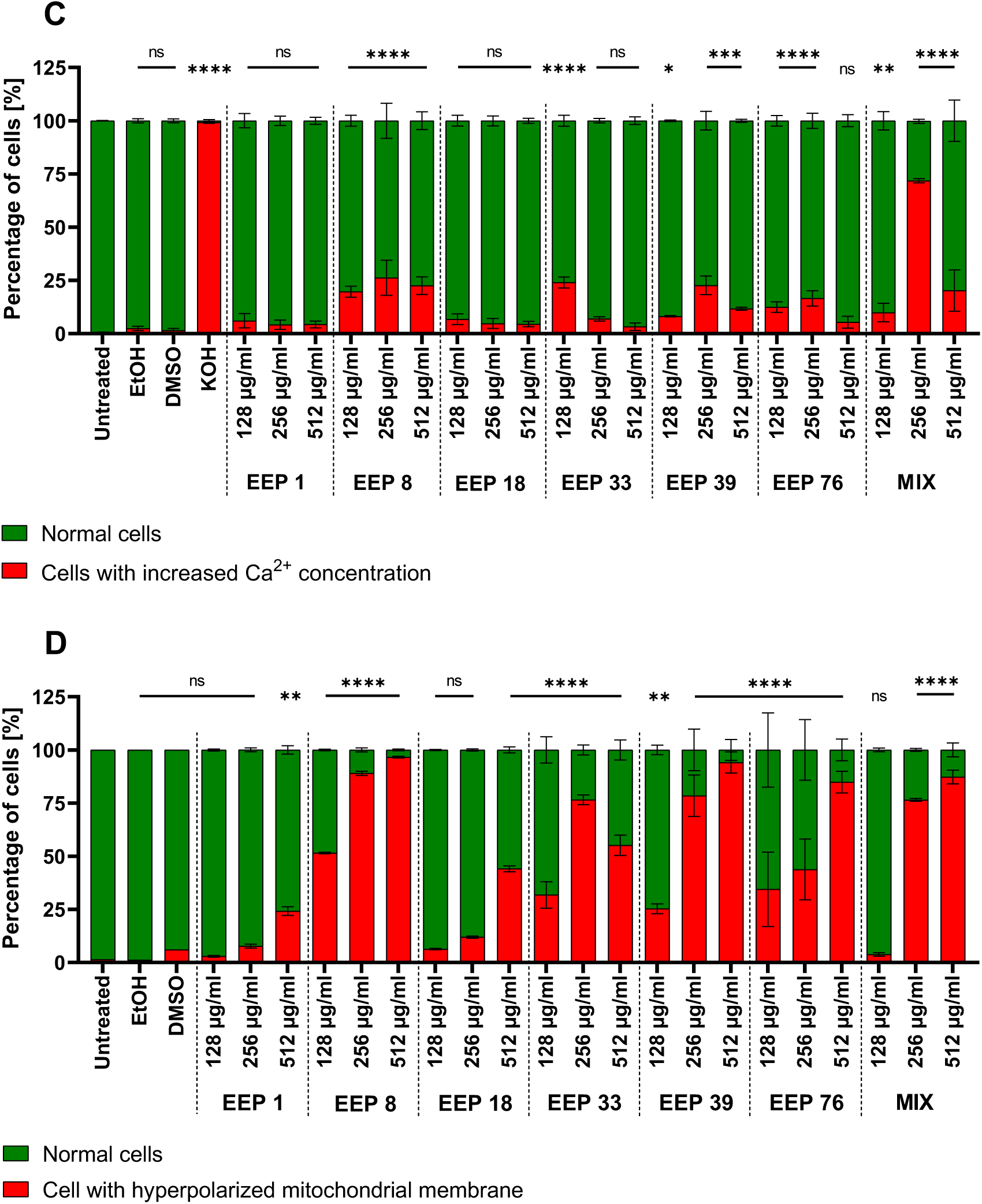
Results of the flow cytometry analyses of *C. albicans* SC 5314 cells exposed to different EEP or CIEEP mixture in concentrations of 128, 256, and 512 µg/mL using different fluorescent dyes: A – PI; B – H_2_-DCFDA; C – Fluo-3-AM; D – Rh123. Statistical significance estimated for fluorescent shift-positive cells, where: *P <0.05, **P <0.01, ***P<0.001, and ****P<0.0001.

#### 3.3.2. Oxidative stress induction

The analysis demonstrated that all samples induced an increase in ROS-positive *C. albicans* cells after 3 h, with extract-specific patterns and predominantly concentration-dependent responses (Fig. 2B). ROS levels in untreated cells were low and comparable to solvent controls (ethanol and DMSO), whereas 1% H₂O₂ produced a strong shift toward ROS-positive cells (47.02%), confirming assay responsiveness. Among the tested extracts, EEP 33 produced consistently high ROS-positive populations already at 128 µg/mL (55.82%) and remained elevated at 256-512 µg/mL (56.47-57.73%), with all concentrations statistically significant versus the control. EEP 39 and EEP 8 also induced statistically significant and robust ROS accumulation, with increasing trends at higher concentrations. EEP 76 increased ROS-positive cells at 128 and 256 µg/mL (44.78% and 60.02%, respectively), but showed a lower fraction at 512 µg/mL (46.8%), despite remaining statistically significant, indicating a non-monotonic response. In contrast, EEP 1 and EEP 18 exhibited weaker and more variable responses at 128 µg/mL that were not statistically significant, but both became significant at 256-512 µg/mL, with moderate ROS-positive populations. The CIEEP mixture induced a graded and statistically significant increase in ROS-positive cells across all concentrations. Although several extracts showed a general tendency toward increased ROS levels with increasing concentration, the observed plateaus or decreases at 512 µg/mL (e.g. EEP 76) may result from loss of measurable signal in severely damaged or rapidly killed cells. This may be due to reduced esterase activity required for H₂-DCFDA conversion or decreased recovery of severely affected cells during cytometric acquisition (Ameziane-El-Hassani & Dupuy, 2013), which is consistent with Time-kill assay results. The strong ROS induction observed for the most active extracts is consistent with previous reports linking propolis exposure to oxidative damage in fungi (Silva-Beltrán et al., 2023). Excess intracellular ROS promotes membrane damage by initiating lipid peroxidation of plasma membrane lipids, which leads to oxidative chain reactions that compromise membrane integrity and permeability (Shahina et al., 2022; Swenson et al., 2024). Free radical production and oxidative injury observed in EEP-treated cells indicate ROS accumulation as an important factor in the early cellular response of *C. albicans* and other yeasts (Madeo et al., 2009; Okińczyc et al., 2020; Yardımcı et al., 2025).

#### 3.3.3. Changes in cytosolic Ca^2+^ level

In addition to ROS, Ca²⁺ ions play a significant role in the initiation and execution of yeast apoptosis (Carraro & Bernardi, 2016). Increased cytosolic Ca²⁺ can lead to mitochondrial permeabilization and the release of proapoptotic factors, thereby initiating programmed cell death. Analysis using Fluo-3 AM (Fig. 2C) after 3 h of exposure revealed that untreated *C. albicans* cells and solvent controls (ethanol, DMSO) consisted almost exclusively of Fluo-3-negative cells, confirming that the assay conditions and solvents did not affect cytosolic Ca²⁺ homeostasis. Addition of KOH produced a result consistent with the well-established ability of alkaline and ionic stresses to induce robust Ca²⁺ influx and activate calcineurin-dependent stress pathways in *C. albicans* (Sanglard et al., 2021; Giuraniuc et al., 2023).

Across the EEP panel, extracts 1 and 18 did not significantly increase the proportion of Fluo-3-positive cells at any tested concentration, with positive events generally remaining below 11% and statistically indistinguishable from the control. In contrast, several extracts exhibited marked and concentration-dependent increases in Fluo-3-positive cells. EEP 8 and EEP 33 induced significant Ca²⁺ elevation already at 128 µg/mL (approximately 19.7% of cells), with EEP 8 showing further enhancement at 256 µg/mL followed by a slight decrease at 512 µg/mL, while EEP 33 exhibited a strong response at 128 µg/mL that decreased at higher concentrations. EEP 39 and EEP 76 also increased the proportion of Fluo-3-positive cells at 128 and 256 µg/mL (approximately 8-29% positive cells), with statistically significant differences compared to the control for all conditions except the highest concentration of EEP 76, where the proportion of Fluo-3-positive cells returned to near-baseline levels. The CIEEP mixture produced one of the most pronounced effects, particularly at 256 µg/mL, where approximately 71.8% of cells were Fluo-3-positive, approaching the level observed under alkaline stress and indicating a strong disturbance of cytosolic Ca²⁺ homeostasis. At 512 µg/mL, the CIEEP mixture also significantly increased the proportion of Ca²⁺-loaded cells, although to a lesser extent than at 256 µg/mL. Overall, these data indicate that a subset of EEPs (notably EEP 8, EEP 33, EEP 39, EEP 76, and especially the CIEEP mixture) induces a notable shift in *C. albicans* populations toward a high-Ca²⁺ state, while others remain largely inactive with respect to Ca²⁺ influx, highlighting extract-specific properties. The observation of distinct maxima at specific concentrations for the most active samples (256 µg/mL for EEP 8, 39, 76, and MIX, or 128 µg/mL for EEP 33) may suggest a threshold for the transition from apoptosis-like responses to necrosis.

To the best of our knowledge, there are no prior reports directly linking propolis treatment to alterations in cytosolic Ca²⁺ levels in yeast, and this study is the first to demonstrate EEP-induced perturbation of Ca²⁺ homeostasis in *C. albicans*. The observed increases in Fluo-3-positive cells are consistent with previous reports in which fungicidal agents induced sustained elevation of cytosolic Ca²⁺ in *C. albicans*, often as part of an apoptosis-like or severe stress response (Sun et al., 2022). For example, carvacrol and related natural products significantly increase the proportion of Fluo-3-positive *C. albicans* cells compared with untreated controls, and this Ca²⁺ overload has been linked to mitochondrial dysfunction and apoptotic cell death (Niu et al., 2020). Similarly, Jia et al. (2019) demonstrated that coumarin induces elevation of cytosolic Ca²⁺ in *C. albicans*, leading to mitochondrial overload, cytochrome c release, and apoptosis. The strong Ca²⁺ influx signatures detected here for EEP 8, EEP 33, EEP 39, EEP 76, and particularly the CIEEP mixture therefore suggest that these propolis extracts may exert part of their antifungal effect through disruption of Ca²⁺ homeostasis and engagement of the Ca²⁺-calcineurin axis, consistent with the recognized role of this pathway in stress adaptation, drug tolerance, and virulence in *C. albicans* (Reedy et al., 2010; Liu et al., 2015; Li et al., 2021).

#### 3.3.4. Hyperpolarization of mitochondrial membrane

Flow cytometry analysis using Rh123 staining demonstrated that EEP induces mitochondrial membrane hyperpolarization in *C. albicans* after 3 h of exposure, with extract- and dose-dependent effects (Fig. 2D). The untreated control showed only 1.38% hyperpolarized cells, while solvent controls (EtOH and DMSO) did not exhibit significant shifts (1.2% and 5.97%, respectively). The EEP-induced shifts toward higher Rh123 fluorescence (hyperpolarization) were validated using positive (oligomycin A, hyperpolarizing) and negative (CCCP, depolarizing) controls, confirming the sensitivity of the assay to mitochondrial membrane potential changes (Figure S2). The most active extracts (EEP 8, 33, 39, and 76) induced statistically significant hyperpolarization at all tested concentrations (128-512 µg/mL), with EEP 8 showing the strongest effect (51.6% at 128 µg/mL, 89% at 256 µg/mL, and 96.6% at 512 µg/mL). EEP 1 and EEP 18 induced significant hyperpolarization only at 512 µg/mL (24.2% and 44.1%), while lower concentrations remained non-significant (≤6.3%). The CIEEP mixture exhibited a concentration-dependent effect, being non-significant at 128 µg/mL (3.8%) but significant at 256 µg/mL (76.5%) and 512 µg/mL (87.3%).

Experimental evidence suggests that mitochondrial membrane hyperpolarization may serve as an early indicator of mitochondrial dysfunction, frequently preceding a cascade of lethal cellular events. For example, treatment with dill seed essential oil (Chen et al., 2013), cannabidiol (Feldman et al., 2021), and peptides (such as C18) (Sun et al., 2022) has been shown to induce significant hyperpolarization, followed by increased mitochondrial ROS production and a decrease in intracellular ATP levels. This bioenergetic disruption is associated with fungicidal activity, as the resulting oxidative stress may lead to chromatin condensation and activation of programmed cell death pathways (Lopes, 2014; Sun et al., 2022).

#### 3.3.5. Hyphae/Pseudohyphae formation

All analyzed EEPs effectively suppressed the morphological transition from blastospores to hyphal or pseudohyphal forms in *C. albicans* after 3 h of exposure (Fig. 3). The proportion of blastospore-sized cells remained high across all treated samples, indicating inhibition of hyphal elongation. In the untreated control (3 h), hyphae/pseudohyphae constituted 65.2% of the population, reflecting spontaneous hyphal induction under the assay conditions. Solvent controls also showed a statistically significant increase in filamentation (EtOH: 38.2%; DMSO: 61.9%).

**Figure 3.**
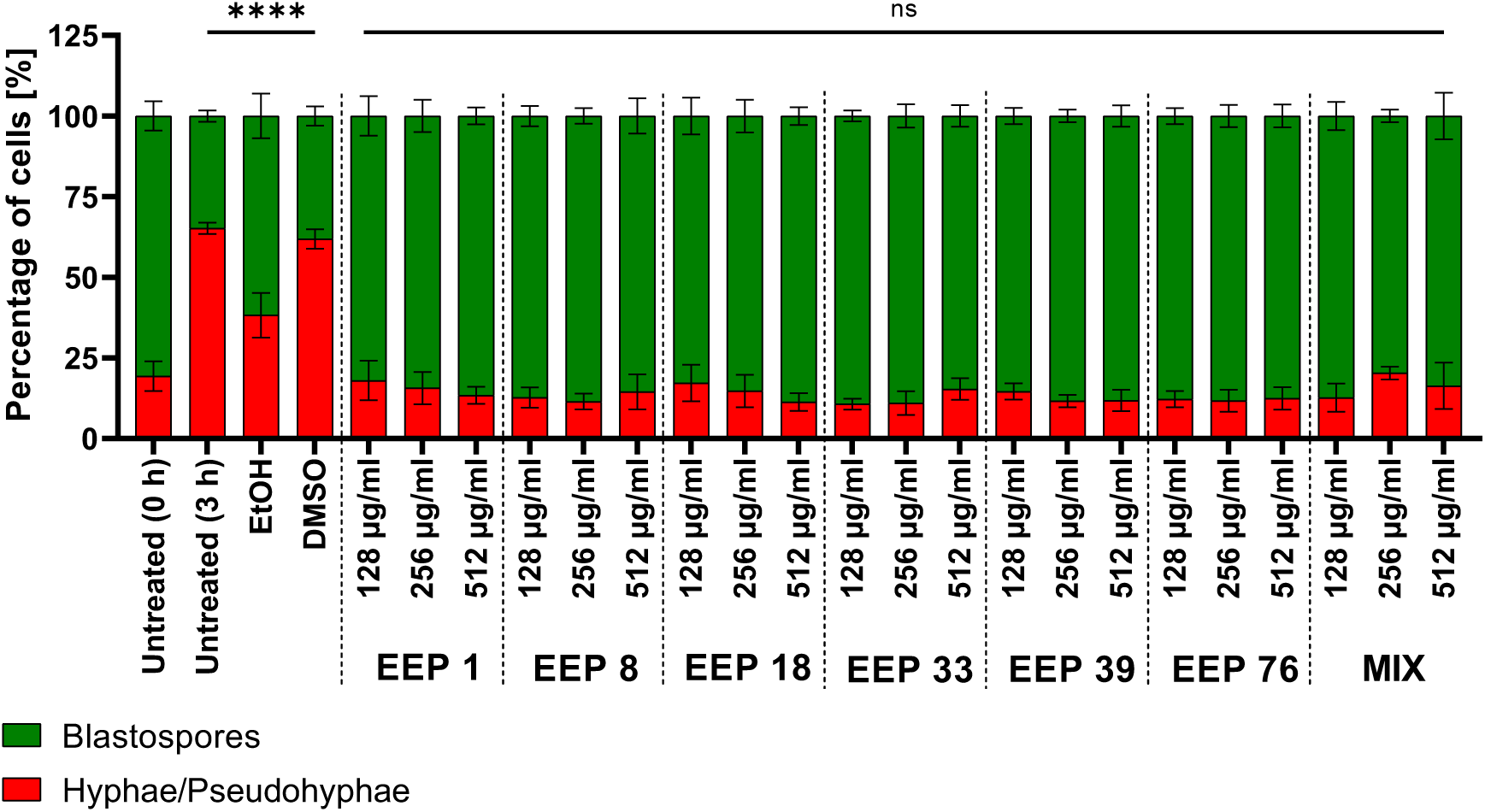
Result of the cytometric analyses of *C. albicans* SC 5314 cell morphological forms adopted under exposure to different EEP or CIEEP mixture in concentrations of 128, 256, and 512 µg/mL. Statistical significance estimated for hyphae/pseudohyphae-positive cells, where: *P <0.05, **P <0.01, ***P<0.001, and ****P<0.0001.

EEP-treated samples consistently maintained high blastospore proportions, with mean values exceeding 82% at all tested concentrations (128, 256, and 512 µg/mL) for EEP 1, EEP 8, EEP 18, EEP 33, EEP 39, and EEP 76. No statistically significant difference in morphology was observed between EEP-treated samples and the untreated control (0 h), suggesting that the extracts prevented the time-dependent increase in filamentation observed in untreated and solvent controls. This inhibitory effect on dimorphism is consistent with propolis-mediated disruption of cell membrane integrity, which interferes with hyphal elongation and germ tube formation (D’Auria et al., 2003; Corrêa et al., 2020). Propolis components have been shown to reduce hyphal formation by damaging the plasma membrane, thereby limiting the morphological plasticity essential for *C. albicans* virulence (Corrêa et al., 2020). By preventing these morphological changes, the extracts counteract the hyphal transition associated with enhanced tissue invasion and biofilm persistence (Haghdoost et al., 2016; Corrêa et al., 2020). These findings support the hypothesis that propolis targets membrane integrity as part of a broader antifungal mechanism, affecting both proliferation and morphological differentiation.

### 3.4. Observation of the effect of EEP/CIEEP mixture on yeast cells using FM

To observe possible morphological changes in *C. albicans* cells under the influence of EEP or the CIEEP mixture, as well as to visually verify the results obtained from MIC/MFC assays with supplemented media and flow cytometry, co-staining with calcofluor white and propidium iodide, as well as co-staining of propidium iodide with MitoTracker™ Green, were performed. Observations were carried out using fluorescence microscopy. Calcofluor white (CFW) is a non-specific chemifluorescent blue dye, particularly effective for visualizing fungal morphology and identifying cell wall structures (Harrington & Hageage, 2003). It binds to β-linked polysaccharides (specifically β-1,3 and β-1,4 polymers) present in chitin and cellulose. Because the *C. albicans* cell wall is rich in chitin, especially at hyphal tips and bud scars, CFW causes these structures to fluoresce brightly under blue light (Watanabe et al., 2005). MitoTracker™ Green (MTG), in contrast, is a cell-permeant fluorescent dye used to visualize mitochondria in living cells. It passively diffuses across the cell membrane and accumulates in the mitochondrial environment, where its mildly reactive chloromethyl group enables covalent binding to thiol groups of mitochondrial proteins (Neikirk et al., 2023).

Co-staining with calcofluor white and PI revealed no visible changes in cell wall integrity and morphology in either control or EEP- or CIEEP-treated samples (Fig. 4A). Control samples were generally characterized by a higher cell density, with only a small fraction of PI-positive cells, indicating intact plasma membranes. A notable observation in both EtOH and DMSO controls was the presence of numerous elongated cells consistent with pseudohyphae or hyphal germ tubes (Sudbery et al., 2004), clearly visible in images acquired at 20× magnification (Fig. S3). In cultures treated with EEP or the CIEEP mixture, the overall cell density was markedly reduced, and most of the visible cells were PI-positive, indicating membrane damage and confirming the strong fungicidal activity of the tested compounds. Notably, round blastospore forms predominated in these samples. Similar observations regarding cell density and morphology were made following co-staining with PI and MitoTracker™ Green (Fig. 4B). Importantly, only cells in control samples exhibited green fluorescence corresponding to stained mitochondria (GFP channel), whereas treated samples showed no detectable mitochondrial signal. This may indicate loss of mitochondrial function following membrane disruption. Additional FM images acquired at 20× and 100× magnification are provided in the Supplementary Material (Figs. S3-S6).

**Figure 4.**
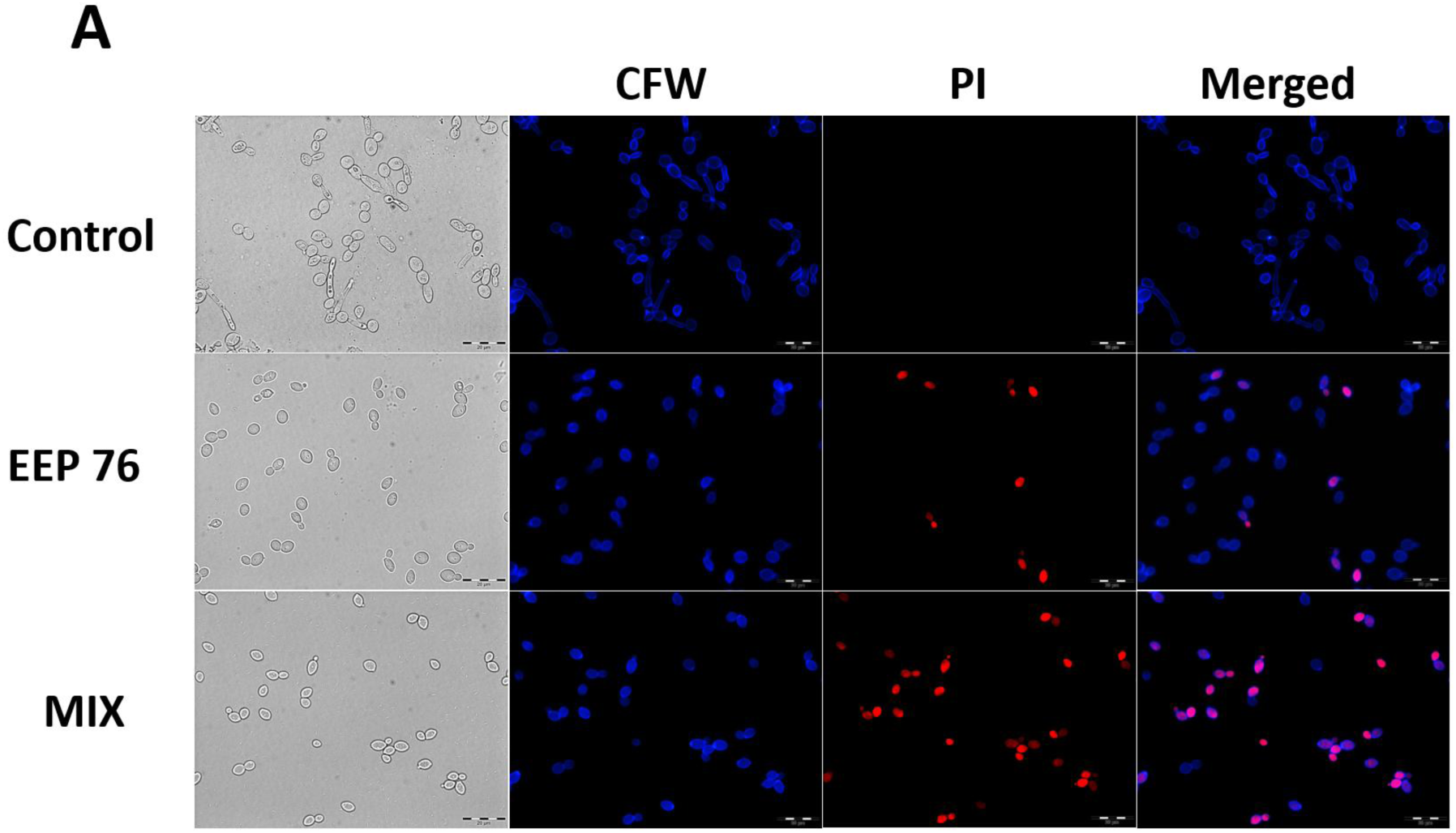
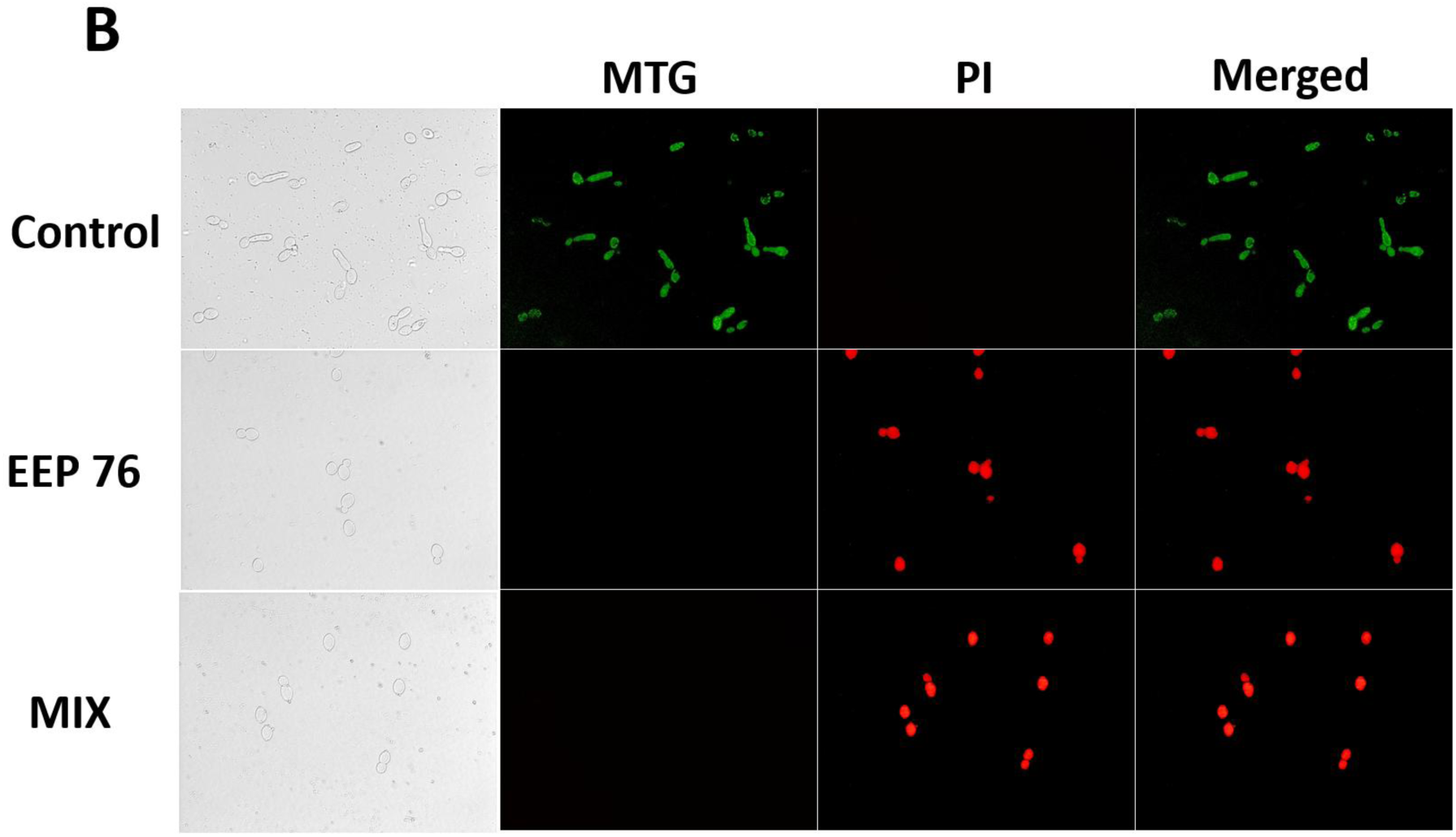
Bright-field and fluorescent microscopy pictures (60x magnification) of *C. albicans* cells treated with EtOH/DMSO (control), EEP 76, and MIX co-stained using: A – Calcofluor White (CFW) and propidium iodide (PI) dyes; B – Mitotracker™ Green (MTG) and propidium iodide (PI) dyes.

### 3.5. EEP induces multi-pathway transcriptional reprogramming in *C. albicans*

RNA-seq data quality and overall sample relationships were first assessed using principal component analysis (PCA) and sample-to-sample distance clustering. PCA based on rlog-transformed expression values revealed clear separation between control and EEP 76-treated samples, with tight clustering of biological replicates (Fig. S7). This pattern was further supported by hierarchical clustering of Euclidean distances, confirming high similarity within experimental groups and distinct separation between conditions (Fig. S8). These results indicate good reproducibility and the absence of major outliers, supporting the validity of downstream differential expression analysis. A comprehensive list of differentially expressed genes identified by DESeq2 is provided in the Supplementary Materials (SCP_vs_SCE_deseq2_annotated.xlsx), including all genes regardless of adjusted p-value. While downstream analyses in this study were primarily based on a more stringent cutoff of |log₂FC| > 2 and adjusted p-value < 0.05, less restrictive thresholds (e.g., |log₂FC| > 1) and unfiltered datasets were also examined to evaluate the robustness of functional enrichment results and to avoid potential bias introduced by arbitrary filtering criteria.

Transcriptomic analysis of *Candida albicans* SC 5314 exposed to EEP 76 revealed 358 differentially expressed genes (DEGs) meeting strict significance criteria (cutoff of |log₂FC| > 2; adjusted p-value < 0.05). Of these, 255 genes were significantly upregulated, including 246 protein-coding genes and 9 non-coding RNAs (5 ncRNAs, 3 rRNAs, and 1 snRNA), with 200 encoding proteins of known biological function. In contrast, all 103 significantly downregulated transcripts were protein-coding, of which 63 had annotated functions (UniProt, 2025). A heatmap of the top 50 most significantly differentially expressed genes, based on Z-scores of rlog-transformed expression values, is presented in Fig. 5.

**Figure 5.**
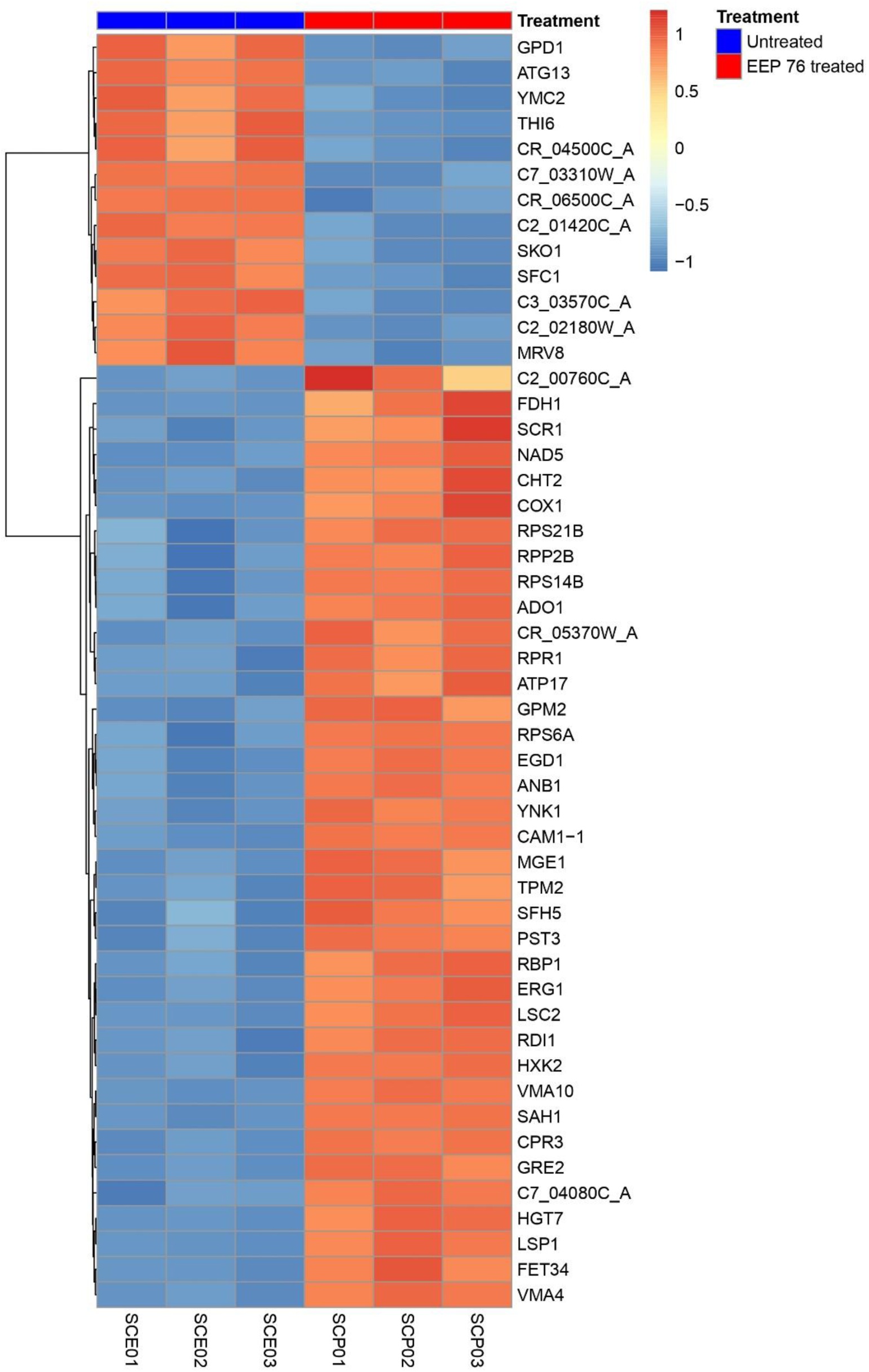
Heatmap of the top-most significantly differentially expressed genes in *C. albicans* SC 5314 following exposure to EEP 76, filtered with log₂FC ≥ 2. Expression values were rlog-transformed and row-wise Z-scored. Samples are ordered by condition: untreated control (SCE01-SCE03) and EEP 76 treated (SCP01-SCP03). Rows (genes) are hierarchically clustered. Gene names were mapped from systematic locus identifiers using Ensembl Fungi BioMart annotations; locus tags are retained where no standard gene name was available.

The identified genes encode proteins involved in virulence, drug response, metabolism, and stress adaptation. In particular, several ATP-binding cassette (ABC) and major facilitator superfamily (MFS) transporters linked to multidrug and azole resistance (CDR1, CDR2, CDR4, CDR11, SNQ2, YOR1, and FLU1), together with the lipid-related transporter RTA3 and the glucose transporter HGT7, were strongly induced, indicating activation of a broad drug response that may reduce intracellular xenobiotic accumulation (Ogawa et al., 1998; Calabrese & Sanglard, 2000; Whaley et al., 2016; Whaley et al., 2018; Laurian et al., 2019). Upregulation was also observed for genes associated with cell wall structure and morphogenesis, including the chitinase CHT2, GPI-anchored proteins such as PGA30 and PGA38, regulators including ACE2 and ROM2, and the filamentous growth regulator FGR41 (Kelly et al., 2004; Plaine et al., 2008; Kanno et al., 2015; Chudzik-Rząd et al., 2022). Additionally, EEP exposure induced a stress and protein homeostasis response, characterized by increased expression of multiple molecular chaperones and co-chaperones (HSP21, HSP60, HSP70/SSA1, SSA2, HSP78, HSP90, HSP104, STF2), as well as numerous ribosomal proteins and translation factors, indicating protein folding stress and a compensatory effort to maintain protein synthesis (Mayer et al., 2013; Gong et al., 2017; Iyer et al., 2022; Choudhary et al., 2023; Robbins & Cowen, 2023). Mitochondrial metabolism was among the most affected functional categories. Nearly all mitochondrial electron transport chain components encoded in the mitochondrial genome (COB, COX1, COX2, COX3B, NAD1-NAD6, NAD4L, NAD5, ATP6, ATP9) showed strong induction, suggesting increased respiratory and ATP synthase capacity. This may reflect elevated energy demand associated with stress responses, transporter activity, and cellular repair, or compensation for partial mitochondrial dysfunction (Bambach et al., 2009; Zhang et al., 2018; Liu et al., 2023). Central carbon and redox metabolism were also reprogrammed, as indicated by overexpression of PCK1 (gluconeogenesis), FDH1 (formate dehydrogenase), aldo-keto reductases such as GRE3, ADO1 (adenosine kinase), and other dehydrogenases. These changes suggest increased gluconeogenic flux, utilization of alternative carbon sources, and detoxification of reactive carbonyl compounds (Sakai et al., 1997; Barelle et al., 2006; Williams et al., 2020; Abraham et al., 2022). Upregulation of multiple genes encoding ribosomal proteins (RPL and RPS families) further indicates attempts to preserve translational capacity and adjust biosynthetic pathways under metabolic stress (Li et al., 2015; Qi et al., 2022). In addition, genes involved in metal and oxidative homeostasis, including CUP1 and CUP2 (copper regulation), the ferroxidase FET34, the flavohemoglobin YHB1, and the peroxiredoxin AHP1, were strongly induced. This was accompanied by upregulation of INO1 and other genes related to phospholipid and inositol metabolism, suggesting disturbances in metal balance, membrane composition, and redox homeostasis (Hromatka et al., 2005; Chen et al., 2008; Ziegler et al., 2011; Schwartz et al., 2013). Gene Ontology enrichment analysis of Biological Process terms, performed separately for up- and downregulated gene sets, further supported a clear functional distinction between suppressed and activated pathways (Fig. 6). Downregulated genes were predominantly associated with rRNA processing, DNA repair and replication, and transcription, whereas upregulated genes were enriched in categories related to proteostasis and the ubiquitin-proteasome system, stress response, mitochondrial energy metabolism, and translation.

**Figure 6.**
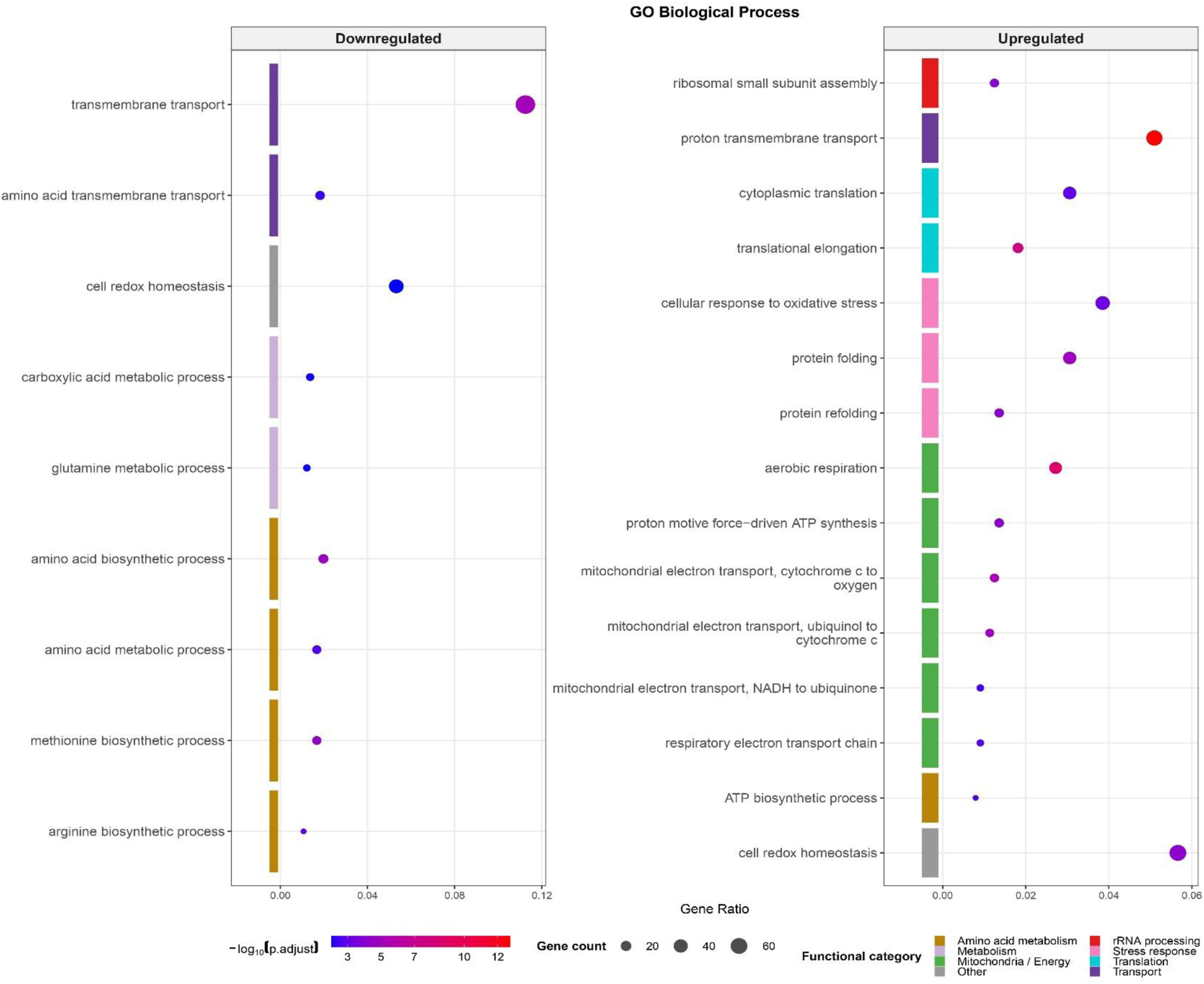
Gene Ontology (GO) Biological Process overrepresentation analysis of differentially expressed genes (log₂FC ≥ 1, adjusted *p*-value < 0.05) in *C. albicans* SC 5314 exposed to EEP 76. The top 15 enriched terms are shown separately for downregulated (left panel) and upregulated (right panel) gene sets. Dot size represents gene count; dot color indicates statistical significance (−log₁₀ adjusted *p*- value). Colored bars along the y-axis denote manually assigned functional categories based on GO term descriptions.

Comparison with the *Saccharomyces cerevisiae* dataset from another study (de Castro et al., 2012) revealed a clear but partial cross-species accordance, with the most consistent overlap observed for genes associated with mitochondrial organization, respiratory metabolism, and stress adaptation. In both species, several genes identified in the *C. albicans* dataset showed concurrent regulation, including mitochondrial and respiration-related genes such as COX8, ATP17, ATP19, and TIM10, as well as stress-associated genes like TRX1, GRE2, and CYB5. In addition, selected genes involved in translational and biosynthetic processes (e.g. RPP1B, RPS28B, RPL21A, and RPL23A) also exhibited consistent regulation, supporting the presence of a conserved core response to propolis, centered on mitochondrial function, redox balance, and metabolic adaptation. At the same time, this conservation was incomplete, as a considerable proportion of shared genes – particularly those involved in transport, cell cycle regulation, and RNA processing – displayed opposite regulation between the two species. As the datasets were generated using chemically distinct propolis samples, namely Polish poplar-type propolis in our *C. albicans* study and Brazilian propolis in the *S. cerevisiae* study, these differences likely reflect a combination of species-specific regulatory programs and differences in composition.

Overall, all the observed transcriptional changes indicate that compounds present in ethanolic extracts of Polish propolis impose combined mitochondrial, oxidative, membrane, and cell wall stress. In response, *C. albicans* upregulates efflux transporters, mitochondrial and metabolic pathways, stress-protective chaperones, and extensive cell surface remodeling – processes closely linked to virulence and antifungal drug response. These findings support the conclusion that propolis does not act through a single dominant mechanism but instead induces a coordinated disruption of cellular homeostasis, consistent with a multi-target mode of action.

### 3.6. Changes in relative expression of selected genes (qPCR)

Genes selected for qPCR analysis were identified based on RNA-seq results. Initially, genes with an adjusted *p*-value greater than 0.05 were excluded. From the remaining set, genes with a log₂FC below 2 (for upregulated genes) or above −2 (for downregulated genes) were further removed. Subsequently, genes annotated as hypothetical or those with only domain-based functional predictions were excluded. This filtering strategy resulted in a more refined list of 176 genes. From this set, 12 genes were selected for qPCR analysis to represent a range of expression changes (both high and moderate), diverse cellular functions, and potential involvement in the response to propolis treatment. In addition, four genes associated with morphological switching in *Candida albicans* (ALS1, ALS3, ALS7, and HWP1) were included based on a prior literature search.

To evaluate the molecular response of *C. albicans* to propolis, expression profiles of selected genes were analysed following exposure to three ethanolic extracts of propolis (EEP 1, EEP 33, EEP 76) with distinct antifungal effectiveness (as evidenced by previous experiments) and a mixture of key components of EEP (CIEEP mixture; MIX). Relative gene expression (log₂RQ) was calculated using the Pfaffl method and is presented in Fig. 7. All tested samples significantly affected the expression of *ERG1*, which encodes squalene epoxidase—a rate-limiting enzyme in the ergosterol biosynthesis pathway and a key determinant of fungal membrane integrity (Pasrija et al., 2005). Suppression of ergosterol biosynthesis emerged as a central response across all treatments. The *ERG1* gene was strongly downregulated in all conditions; however, the highly active samples (EEP 33, EEP 76, and MIX) induced markedly stronger downregulation (log₂RQ ranging from −21.41 to −37.81) compared to the low-activity EEP 1 (−11.53). This disruption of sterol biosynthesis was accompanied by a pronounced upregulation of the lipase gene *LIP3* (log₂RQ > 19.32) across all samples. In parallel, all EEPs triggered a strong, extract-specific upregulation of the secreted aspartyl proteinase gene *SAP4* (up to 63.92 in EEP 76).

**Figure 7.**
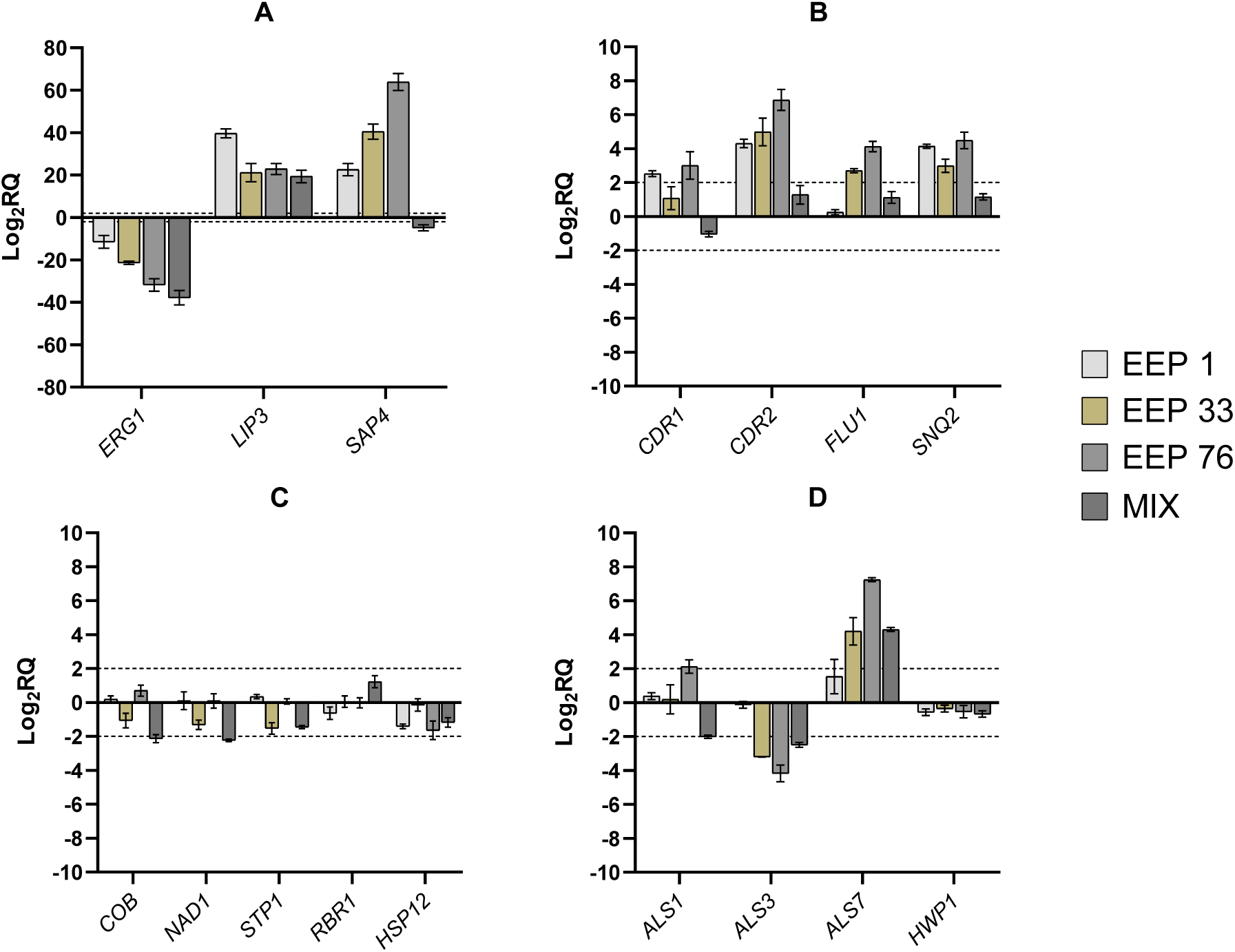
Relative gene expression determined for selected genes of *C. albicans* exposed to 192 µg/mL of EEP 1, EEP 33, EEP 76, and MIX for 3 h, expressed as log_2_RQ. Graphs include the data for following genes: A – *ERG1*, *LIP3*, and *SAP4*; B – *CDR1*, *CDR2*, *FLU1*, and *SNQ2*; C – *COB*, *NAD1*, *STP1*, *RBR1*, and *HSP12*. The values were considered biologically significant at an absolute threshold of log_2_RQ ≥ 2 (for upregulated genes) or log_2_RQ ≤ -2 (for downregulated genes) – indicated by dotted lines.

Because *SAP4* and *LIP3* encode extracellular virulence factors involved in host tissue penetration, protein degradation, and lipid utilization (Naglik et al., 2003; Paraje et al., 2008), their strong induction may represent a stress response aimed at compensating for disrupted membrane biosynthesis. Notably, the upregulation of *SAP4* and *LIP3* contrasts with reports describing propolis as an inhibitor of *Candida* virulence factors (Tobaldini-Valerio et al., 2016). Unlike EEP, the CIEEP mixture reduced *SAP4* expression (log₂RQ = −4.87) and was the only treatment to induce biologically significant downregulation of mitochondrial respiration genes (*COB*, −2.13; *NAD1*, −2.23). This suggests that the defined mixture of key components may exert an additional mechanism targeting fungal energy metabolism that is not fully recapitulated by whole extracts. EEP-treated cells consistently upregulated the ABC transporter genes *CDR2* and *SNQ2*, which mediate multidrug resistance and active efflux (Holmes et al., 2008), indicating activation of xenobiotic export pathways in response to complex extract composition. The multidrug transporters *FLU1* and *CDR1* were additionally upregulated by EEP 1 and EEP 76. In contrast, none of these efflux-related genes exceeded the assumed biological significance threshold following CIEEP mixture treatment. Fungal ABC and MFS transporters are known to be strongly modulated by natural plant-derived compounds, reflecting adaptive responses to xenobiotic stress (Gil et al., 2022). Furthermore, transcriptomic profiling revealed a relationship between antifungal efficacy and the expression of agglutinin-like sequence (ALS) genes, which mediate adhesion and biofilm formation (Hoyer, 2001). The *ALS3* gene, encoding a major hypha-associated adhesin critical for host cell attachment (Liu & Filler, 2011), was significantly downregulated by the CIEEP mixture and by EEP 33 and 76. In contrast, *ALS7* was significantly upregulated by EEP 33, EEP 76, and the CIEEP mixture, while *ALS1* was downregulated exclusively by the CIEEP mixture. The simultaneous downregulation of *ALS3* and upregulation of *ALS7* may reflect compensatory transcriptional reprogramming of adhesion-related functions in response to propolis-induced stress. While Als3p serves as a primary multifunctional adhesin, its suppression may be partially offset by induction of *ALS7*, which encodes a structurally related but functionally distinct adhesin (Zhao et al., 2007). The suppression of *ALS3* is consistent with an anti-biofilm effect and supports previous studies demonstrating that propolis and its derivatives reduce *ALS3* expression, thereby inhibiting biofilm formation and morphological transition (Iadnut et al., 2019).

During the comparison of RNA-seq and qPCR results, a discrepancy was observed for *ERG1*, which was upregulated in the RNA-seq dataset but strongly downregulated in qPCR analysis. Similar inconsistencies between RNA-seq and RT-qPCR have been reported previously, where overall agreement between methods is generally high, but a subset of genes displays discordant expression patterns due to gene-specific technical and analytical factors. These include differences in transcript characteristics, quantification approaches, and normalization strategies, as well as platform-specific biases in sensitivity and dynamic range (Sun et al., 2014; Everaert et al., 2017). In this context, the opposing regulation of *ERG1* is unlikely to reflect a global inhibition of ergosterol biosynthesis, particularly given that multiple key genes within the pathway (*ERG1, ERG6, ERG9, ERG20, ERG26,* and *ERG28*) were consistently upregulated in the RNA-seq data. Instead, this pattern may indicate a complex, compensatory response of sterol metabolism to propolis-induced membrane stress, in which pathway-level activation coexists with gene-specific regulatory divergence.

## 4. Conclusions

This study provides a comprehensive mechanistic insight into the antifungal activity of ethanolic extracts of Polish propolis against *Candida albicans*. The obtained results consistently demonstrate that propolis exerts a potent, concentration-dependent antifungal effect characterized by rapid fungicidal activity in the most active samples. Mechanistic investigations indicate that the primary site of action is the fungal cell membrane, as supported by increased membrane permeability and reduced efficacy in the presence of exogenous ergosterol. At the cellular level, propolis exposure induces a cascade of intracellular disturbances, including reactive oxygen species accumulation, mitochondrial dysfunction, disruption of Ca²⁺ homeostasis, and inhibition of hyphal formation, collectively impairing both viability and virulence of *C. albicans*. Importantly, transcriptomic analysis revealed that these effects are accompanied by a global reprogramming of gene expression, involving downregulation of essential growth-related processes and concurrent activation of stress-response pathways. These findings indicate that propolis does not act through a single molecular target but rather induces a coordinated, multi-level disruption of fungal cellular homeostasis. Taken together, this study supports the concept of propolis as a multi-target antifungal agent with both fungicidal and antivirulence properties. Given its complex composition and broad mechanism of action, propolis represents a promising candidate for the development of alternative or adjunct antifungal strategies. Further studies should focus on standardization of active components, preparing effective formulations, *in vivo* validation of these formulations (mostly for treatment of topical fungal infections), and evaluation of potential synergistic interactions with conventional antifungal agents. A better understanding of the molecular mechanism of the biological activity of propolis is necessary for more widespread and more rational application of this product preparations in modern medicine.

## CRediT authorship contribution statement

**Piotr Bollin:** Writing – review & editing, Writing – original draft, Methodology, Investigation, Data curation, Conceptualization. **Michał K. Pierański:** Writing – review & editing, Writing – original draft, Project administration, Methodology, Investigation, Data curation, Conceptualization. **Michał W. Szcześniak:** Writing – review & editing, Writing – original draft, Software, Methodology, Investigation, Data curation. **Piotr Szweda:** Writing – review & editing, Supervision, Project administration, Methodology, Funding acquisition, Formal analysis, Data curation, Conceptualization.

## Data availability

The RNA sequencing data generated in this study have been deposited in the Gene Expression Omnibus (GEO) under accession number (GSE328537). All other data supporting the findings of this study are included in the article and its Supplementary Materials. Additional raw data are available from the corresponding authors upon reasonable request.

## Declaration of generative AI and AI-assisted technologies in the manuscript preparation process

During the preparation of this work the authors used ChatGPT (OpenAI, GPT-5.3 model, https://openai.com/chatgpt) in order to to support language editing and stylistic refinement of the manuscript. After using this tool, the authors reviewed and edited the content as needed and take full responsibility for the content of the published article.

## Abbreviations

BSA: bovine serum albumin
CCCP: carbonyl cyanide m-chlorophenyl hydrazone
CFW: Calcofluor White
CIEEP: crucial ingredients of ethanolic extract of propolis
CLSI: Clinical Laboratory Standard Institute
DMSO: dimethyl sulfoxide
EEP: ethanolic extract of propolis
FC: fold change
FCM: flow cytometry
FSC-H: forward scatter height
H_2_-DCFDA: 2′,7′-dichlorodihydrofluorescein diacetate
HEPES: 4-(2-hydroxyethyl)-1-piperazineethanesulfonic acid
MFC: minimum fungicidal concentration
MIC: minimum inhibitory concentration
MOPS: 3-N-morpholinopropanesulfonic acid
MTG: Mitotracker™ Green
PBS: phosphate-buffered saline
PI: propidium iodide
RT-qPCR: quantitative reverse transcription PCR
qPCR: quantitative PCR
Rh123: rhodamine 123
ROS: reactive oxygen species
RT: reverse transcription
SD: standard deviation
RNA-seq: RNA sequencing
YPD: Yeast Extract Peptone Dextrose

## References

1. Abraham, N., Schroeter, K. L., Zhu, Y., Chan, J., Evans, N., Kimber, M. S., Carere, J., Zhou, T., & Seah, S. Y. K. (2022). Structure-function characterization of an aldo-keto reductase involved in detoxification of the mycotoxin, deoxynivalenol. Scientific Reports, 12(1), 14737. 10.1038/s41598-022-19040-8

2. Ameziane-El-Hassani, R. and Dupuy, C. (2013). Detection of Intracellular Reactive Oxygen Species (CM-H_2_DCFDA). Bio-protocol, 3(1): e313. 10.21769/BioProtoc.313

3. Andrews, S. (2010). FastQC: a quality control tool for high throughput sequence data. Available online at: https://www.bioinformatics.babraham.ac.uk/projects/fastqc/

4. Ansari, J. A., Naz, S., Tarar, O. M., Siddiqi, R., Haider, M. S., & Jamil, K. (2015). Binding effect of proline-rich-proteins (PRPs) on *in vitro* antimicrobial activity of the flavonoids. Brazilian Journal of Microbiology, 46(1), 183–188. 10.1590/S1517-838246120130280

5. Bambach, A., Fernandes, M. P., Ghosh, A., Kruppa, M., Alex, D., Li, D., Fonzi, W. A., Chauhan, N., Sun, N., Agrellos, O. A., Vercesi, A. E., Rolfes, R. J., & Calderone, R. (2009). Goa1p of *Candida albicans* localizes to the mitochondria during stress and is required for mitochondrial function and virulence. Eukaryotic Cell, 8(11), 1706–1720. 10.1128/EC.00066-09

6. Barelle, C. J., Priest, C. L., Maccallum, D. M., Gow, N. A., Odds, F. C., & Brown, A. J. (2006). Niche-specific regulation of central metabolic pathways in a fungal pathogen. Cellular Microbiology, 8(6), 961–971. 10.1111/j.1462-5822.2005.00676.x

7. Bhattacharya, S., Sae-Tia, S., & Fries, B. C. (2020). Candidiasis and Mechanisms of Antifungal Resistance. Antibiotics, 9(6), 312. 10.3390/antibiotics9060312

8. Bollin, P., Kuś, P. M., Okińczyc, P., Van Dijck, P., & Szweda, P. (2025). Identification of potential markers of elevated anticandidal activity of propolis extracts. Journal of Ethnopharmacology, 347, 119799. 10.1016/j.jep.2025.119799

9. Bushnell, B. (2014). BBTools software package. Available online at: https://sourceforge.net/projects/bbmap/

10. Calabrese, D., Bille, J., & Sanglard, D. (2000). A novel multidrug efflux transporter gene of the major facilitator superfamily from *Candida albicans* (FLU1) conferring resistance to fluconazole. Microbiology, 146(11), 2743–2754. 10.1099/00221287-146-11-2743

11. Carraro, M., & Bernardi, P. (2016). Calcium and reactive oxygen species in regulation of the mitochondrial permeability transition and of programmed cell death in yeast. Cell Calcium, 60(2), 102–107. 10.1016/j.ceca.2016.03.005

12. Cerqueira, P., Cunha, A., & Almeida-Aguiar, C. (2022). Potential of propolis antifungal activity for clinical applications. Journal of Applied Microbiology, 133(3), 1207–1228. 10.1111/jam.15628

13. Chen, Y. L., Kauffman, S., & Reynolds, T. B. (2008). *Candida albicans* uses multiple mechanisms to acquire the essential metabolite inositol during infection. Infection and Immunity, 76(6), 2793–2801. 10.1128/IAI.01514-07

14. Chen, Y., Zeng, H., Tian, J., Ban, X., Ma, B., & Wang, Y. (2013). Antifungal mechanism of essential oil from Anethum graveolens seeds against *Candida albicans*. Journal of Medical Microbiology, 62(8), 1175–1183. 10.1099/jmm.0.055467-0

15. Choudhary, S., Mundodi, V., Smith, A. D., & Kadosh, D. (2023). Genome-wide translational response of *Candida albicans* to fluconazole treatment. Microbiology Spectrum, 11(5). 10.1128/spectrum.02572-23

16. Chudzik-Rząd, B., Zalewski, D., Kasela, M., Sawicki, R., Szymańska, J., Bogucka-Kocka, A., & Malm, A. (2022). The Landscape of Gene Expression during Hyperfilamentous Biofilm Development in Oral *Candida albicans* Isolated from a Lung Cancer Patient. International Journal of Molecular Sciences, 24(1), 368. 10.3390/ijms24010368

17. Corrêa, J. L., Veiga, F. F., Jarros, I. C., Costa, M. I., Castilho, P. F., de Oliveira, K. M. P., Rosseto, H. C., Bruschi, M. L., Svidzinski, T. I. E., & Negri, M. (2020). Propolis extract has bioactivity on the wall and cell membrane of *Candida albicans*. Journal of Ethnopharmacology, 256, 112791. 10.1016/j.jep.2020.112791

18. Crowley, L. C., Scott, A. P., Marfell, B. J., Boughaba, J. A., Chojnowski, G., & Waterhouse, N. J. (2016). Measuring Cell Death by Propidium Iodide Uptake and Flow Cytometry. Cold Spring Harbor Protocols, 2016(7), pdb.prot087163. 10.1101/pdb.prot087163

19. Cunningham, F., Allen, J. E., Allen, J., Alvarez-Jarreta, J., Amode, M. R., Armean, I. M., Austine-Orimoloye, O., Azov, A. G., Barnes, I., Bennett, R., Berry, A., Bhai, J., Bignell, A., Billis, K., Boddu, S., Brooks, L., Charkhchi, M., Cummins, C., Da Rin Fioretto, L., … Flicek, P. (2021). Ensembl 2022. Nucleic Acids Research, 50(D1), D988–D995. 10.1093/nar/gkab1049

20. D’Auria, F. D., Tecca, M., Scazzocchio, F., Renzini, V., & Strippoli, V. (2003). Effect of Propolis on Virulence Factors of *Candida albicans*. Journal of Chemotherapy, 15(5), 454–460. 10.1179/joc.2003.15.5.454

21. de Castro, P. A., Savoldi, M., Bonatto, D., Malavazi, I., Goldman, M. H. S., Berretta, A. A., & Goldman, G. H. (2012). Transcriptional profiling of *Saccharomyces cerevisiae* exposed to propolis. BMC Complementary and Alternative Medicine, 12(1). 10.1186/1472-6882-12-194

22. Denning, D. W. (2024). Global incidence and mortality of severe fungal disease. The Lancet Infectious Diseases, 24(7), e428–e438. 10.1016/s1473-3099(23)00692-8

23. Everaert, C., Luypaert, M., Maag, J. L. V., Cheng, Q. X., Dinger, M. E., Hellemans, J., & Mestdagh, P. (2017). Benchmarking of RNA-sequencing analysis workflows using whole-transcriptome RT-qPCR expression data. Scientific Reports, 7(1). 10.1038/s41598-017-01617-3

24. Ewels, P., Magnusson, M., Lundin, S., & Käller, M. (2016). MultiQC: summarize analysis results for multiple tools and samples in a single report. Bioinformatics, 32(19), 3047–3048. 10.1093/bioinformatics/btw354

25. Fabris, S., Bertelle, M., Astafyeva, O., Gregoris, E., Zangrando, R., Gambaro, A., Lima, G. P. P., & Stevanato, R. (2013). Antioxidant Properties and Chemical Composition Relationship of Europeans and Brazilians Propolis. Pharmacology & Pharmacy, 04(01), 46–51. 10.4236/pp.2013.41006

26. Feldman, M., Sionov, R. V., Mechoulam, R., & Steinberg, D. (2021). Anti-Biofilm Activity of Cannabidiol against *Candida albicans*. Microorganisms, 9(2), 441. 10.3390/microorganisms9020441

27. Gil, F., Laiolo, J., Bayona-Pacheco, B., Cannon, R. D., Ferreira-Pereira, A., & Carpinella, M. C. (2022). Extracts from Argentinian native plants reverse fluconazole resistance in *Candida* species by inhibiting the efflux transporters Mdr1 and Cdr1. BMC Complementary Medicine and Therapies, 22(1), 264. 10.1186/s12906-022-03745-4

28. Giuraniuc, C. V., Parkin, C., Almeida, M. C., Fricker, M., Shadmani, P., Nye, S., Wehmeier, S., Chawla, S., Bedekovic, T., Lehtovirta-Morley, L., Richards, D. M., Gow, N. A., & Brand, A. C. (2023). Dynamic calcium-mediated stress response and recovery signatures in the fungal pathogen, *Candida albicans*. mBio, 14(5), e0115723. 10.1128/mbio.01157-23

29. Gong, Y., Li, T., Yu, C., & Sun, S. (2017). *Candida albicans* Heat Shock Proteins and Hsps-Associated Signaling Pathways as Potential Antifungal Targets. Frontiers in Cellular and Infection Microbiology, 7. 10.3389/fcimb.2017.00520

30. Gucwa, K., Kusznierewicz, B., Milewski, S., Van Dijck, P., & Szweda, P. (2018). Antifungal Activity and Synergism with Azoles of Polish Propolis. Pathogens, 7(2), 56. 10.3390/pathogens7020056

31. Haghdoost, N. S., Salehi, T. Z., Khosravi, A., & Sharifzadeh, A. (2016). Antifungal activity and influence of propolis against germ tube formation as a critical virulence attribute by clinical isolates of *Candida albicans*. Journal de Mycologie Medicale, 26(4), 298–305. 10.1016/j.mycmed.2015.11.004

32. Harrington, B. J., & Hageage, G. J., Jr. (2003). Calcofluor White: A Review of its Uses and Applications in Clinical Mycology and Parasitology. Laboratory Medicine, 34(5), 361–367. 10.1309/eph2tdt8335gh0r3

33. Holmes, A. R., Lin, Y. H., Niimi, K., Lamping, E., Keniya, M., Niimi, M., Tanabe, K., Monk, B. C., & Cannon, R. D. (2008). ABC transporter Cdr1p contributes more than Cdr2p does to fluconazole efflux in fluconazole-resistant *Candida albicans* clinical isolates. Antimicrobial Agents and Chemotherapy, 52(11), 3851–3862. 10.1128/AAC.00463-08

34. Hoyer L. L. (2001). The ALS gene family of *Candida albicans*. Trends in Microbiology, 9(4), 176–180. 10.1016/s0966-842x(01)01984-9

35. Hromatka, B. S., Noble, S. M., & Johnson, A. D. (2005). Transcriptional Response of *Candida albicans* to Nitric Oxide and the Role of theYHB1Gene in Nitrosative Stress and Virulence. Molecular Biology of the Cell, 16(10), 4814–4826. 10.1091/mbc.e05-05-0435

36. Iadnut, A., Mamoon, K., Thammasit, P., Pawichai, S., Tima, S., Preechasuth, K., Kaewkod, T., Tragoolpua, Y., & Tragoolpua, K. (2019). *In Vitro* Antifungal and Antivirulence Activities of Biologically Synthesized Ethanolic Extract of Propolis-Loaded PLGA Nanoparticles against *Candida albicans*. Evidence-Based Complementary and Alternative Medicine, 2019, 1–14. 10.1155/2019/3715481

37. Isidorov, V. A., Bakier, S., Pirożnikow, E., Zambrzycka, M., & Swiecicka, I. (2016). Selective Behaviour of Honeybees in Acquiring European Propolis Plant Precursors. Journal of Chemical Ecology, 42(6), 475–485. 10.1007/s10886-016-0708-9

38. Iyer, K. R., Robbins, N., & Cowen, L. E. (2022). The role of *Candida albicans* stress response pathways in antifungal tolerance and resistance. iScience, 25(3), 103953. 10.1016/j.isci.2022.103953

39. Jia, C., Zhang, J., Yu, L., Wang, C., Yang, Y., Rong, X., Xu, K., & Chu, M. (2019). Antifungal Activity of Coumarin Against *Candida albicans* Is Related to Apoptosis. Frontiers in Cellular and Infection Microbiology, 8. 10.3389/fcimb.2018.00445

40. Kanno, T., Takekawa, D., & Miyakawa, Y. (2015). Analysis of the essentiality of ROM2 genes in the pathogenic yeasts *Candida glabrata* and *Candida albicans* using temperature-sensitive mutants. Journal of Applied Microbiology, 118(4), 851–863. 10.1111/jam.12745

41. Kasote, D., Bankova, V., & Viljoen, A. M. (2022). Propolis: chemical diversity and challenges in quality control. Phytochemistry Reviews, 21(6), 1887–1911. 10.1007/s11101-022-09816-1

42. Kelly, M. T., MacCallum, D. M., Clancy, S. D., Odds, F. C., Brown, A. J. P., & Butler, G. (2004). The *Candida albicans* CaACE2 gene affects morphogenesis, adherence and virulence. Molecular Microbiology, 53(3), 969–983. 10.1111/j.1365-2958.2004.04185.x

43. Kolde, R. (2019). pheatmap: Pretty Heatmaps. R package version 1.0.12. https://CRAN.R-project.org/package=pheatmap

44. Kuropatnicki, A. K., Szliszka, E., & Krol, W. (2013). Historical Aspects of Propolis Research in Modern Times. Evidence-Based Complementary and Alternative Medicine, 2013, 1–11. 10.1155/2013/964149

45. Langmead, B., & Salzberg, S. L. (2012). Fast gapped-read alignment with Bowtie 2. Nature Methods, 9(4), 357–359. 10.1038/nmeth.1923

46. Larkin, E., Hager, C., Chandra, J., Mukherjee, P. K., Retuerto, M., Salem, I., Long, L., Isham, N., Kovanda, L., Borroto-Esoda, K., Wring, S., Angulo, D., & Ghannoum, M. (2017). The Emerging Pathogen *Candida auris*: Growth Phenotype, Virulence Factors, Activity of Antifungals, and Effect of SCY-078, a Novel Glucan Synthesis Inhibitor, on Growth Morphology and Biofilm Formation. Antimicrobial Agents and Chemotherapy, 61(5). 10.1128/aac.02396-16

47. Laurian, R., Dementhon, K., Doumèche, B., Soulard, A., Noel, T., Lemaire, M., & Cotton, P. (2019). Hexokinase and Glucokinases are Essential for Fitness and Virulence in the Pathogenic Yeast *Candida albicans*. Frontiers in Microbiology, 10. 10.3389/fmicb.2019.00327

48. Lee, Y. K., Segars, K. L., & Trinkaus-Randall, V. (2020). Multiple Imaging Modalities for Cell-Cell Communication via Calcium Mobilizations in Corneal Epithelial Cells. Methods in Molecular Biology, 2346, 11–20. 10.1007/7651_2020_329

49. Lee, Y., Puumala, E., Robbins, N., & Cowen, L. E. (2020B). Antifungal Drug Resistance: Molecular Mechanisms in *Candida albicans* and Beyond. Chemical Reviews, 121(6), 3390–3411. 10.1021/acs.chemrev.0c00199

50. Leite, M. C. A., Bezerra, A. P. de B., Sousa, J. P. de, Guerra, F. Q. S., & Lima, E. de O. (2014). Evaluation of Antifungal Activity and Mechanism of Action of Citral against *Candida albicans*. Evidence-Based Complementary and Alternative Medicine, 2014(1). 10.1155/2014/378280

51. Li, B., & Dewey, C. N. (2011). RSEM: accurate transcript quantification from RNA-Seq data with or without a reference genome. BMC Bioinformatics, 12(1). 10.1186/1471-2105-12-323

52. Li, P., Seneviratne, C. J., Alpi, E., Vizcaino, J. A., & Jin, L. (2015). Delicate Metabolic Control and Coordinated Stress Response Critically Determine Antifungal Tolerance of *Candida albicans* Biofilm Persisters. Antimicrobial Agents and Chemotherapy, 59(10), 6101–6112. 10.1128/aac.00543-15

53. Li, W., Shrivastava, M., Lu, H., & Jiang, Y. (2021). Calcium-calcineurin signaling pathway in *Candida albicans*: A potential drug target. Microbiological Research, 249, 126786. 10.1016/j.micres.2021.126786

54. Liu, S., Hou, Y., Liu, W., Lu, C., Wang, W., & Sun, S. (2015). Components of the calcium-calcineurin signaling pathway in fungal cells and their potential as antifungal targets. Eukaryotic Cell, 14(4), 324–334. 10.1128/EC.00271-14

55. Liu, Y., & Filler, S. G. (2011). *Candida albicans* Als3, a multifunctional adhesin and invasin. Eukaryotic Cell, 10(2), 168–173. 10.1128/EC.00279-10

56. Liu, Z., Basso, P., Hossain, S., Liston, S. D., Robbins, N., Whitesell, L., Noble, S. M., & Cowen, L. E. (2023). Multifactor transcriptional control of alternative oxidase induction integrates diverse environmental inputs to enable fungal virulence. Nature Communications, 14(1). 10.1038/s41467-023-40209-w

57. Lopes, G. L. L. (2014). Seaweeds from the Portuguese coast: chemistry, antimicrobial and antiinflammatory capacity [Universidade do Porto]. 10.34626/68NA-8C58

58. Love, M. I., Huber, W., & Anders, S. (2014). Moderated estimation of fold change and dispersion for RNA-seq data with DESeq2. Genome Biology, 15(12). 10.1186/s13059-014-0550-8

59. M27-A2 (2002). Reference Method for Broth Dilution Antifungal Susceptibility Testing of Yeast; Approved Guideline – Second Edition. Clinical and Laboratory Standards Institute (CLSI), Wayne, PA, USA.

60. Madeo, F., Carmona-Gutierrez, D., Ring, J., Büttner, S., Eisenberg, T., & Kroemer, G. (2009). Caspase-dependent and caspase-independent cell death pathways in yeast. Biochemical and Biophysical Research Communications, 382(2), 227–231. 10.1016/j.bbrc.2009.02.117

61. Majorek, K. A., Porebski, P. J., Dayal, A., Zimmerman, M. D., Jablonska, K., Stewart, A. J., Chruszcz, M., & Minor, W. (2012). Structural and immunological characterization of bovine, horse, and rabbit serum albumins. Molecular Immunology, 52(3-4), 174–182. 10.1016/j.molimm.2012.05.011

62. Mayer, F. L., Wilson, D., & Hube, B. (2013). Hsp21 Potentiates Antifungal Drug Tolerance in *Candida albicans*. PLoS ONE, 8(3), e60417. 10.1371/journal.pone.0060417

63. Naglik, J. R., Challacombe, S. J., & Hube, B. (2003). Candida albicansSecreted Aspartyl Proteinases in Virulence and Pathogenesis. Microbiology and Molecular Biology Reviews, 67(3), 400–428. 10.1128/mmbr.67.3.400-428.2003

64. Neikirk, K., Marshall, A. G., Kula, B., Smith, N., LeBlanc, S., & Hinton, A., Jr. (2023). MitoTracker: A useful tool in need of better alternatives. European Journal of Cell Biology, 102(4), 151371. 10.1016/j.ejcb.2023.151371

65. Neuwirth, E. (2022). RColorBrewer: ColorBrewer Palettes. R package version 1.1-3. https://CRAN.R-project.org/package=RColorBrewer

66. Niu, C., Wang, C., Yang, Y., Chen, Y., Zou, W., Liu, H., & Wang, Y. (2020). Carvacrol induces *Candida albicans* apoptosis associated with Ca²⁺/calcineurin pathway. Frontiers in Cell and Developmental Biology, 8, 192. 10.3389/fcell.2020.00192

67. O’Connor, J. E., Vargas, J. L., KimLer, B. F., Hernandez-Yago, J., & Grisolia, S. (1988). Use of Rhodamine 123 to investigate alterations in mitochondrial activity in isolated mouse liver mitochondria. Biochemical and Biophysical Research Communications, 151(1), 568–573. 10.1016/0006-291x(88)90632-8

68. Ogawa, A., Hashida-Okado, T., Endo, M., Yoshioka, H., Tsuruo, T., Takesako, K., & Kato, I. (1998). Role of ABC transporters in aureobasidin A resistance. Antimicrobial Agents and Chemotherapy, 42(4), 755–761. 10.1128/AAC.42.4.755

69. Okińczyc, P., Paluch, E., Franiczek, R., Widelski, J., Wojtanowski, K. K., Mroczek, T., Krzyżanowska, B., Skalicka-Woźniak, K., & Sroka, Z. (2020). Antimicrobial activity of *Apis mellifera L.* and *Trigona sp*. propolis from Nepal and its phytochemical analysis. Biomedicine & Pharmacotherapy, 129, 110435. 10.1016/j.biopha.2020.110435

70. Okińczyc, P., Widelski, J., Ciochoń, M., Paluch, E., Bozhadze, A., Jokhadze, M., Mtvarelishvili, G., Korona-Głowniak, I., Krzyżanowska, B., & Kuś, P. M. (2022). Phytochemical Profile, Plant Precursors and Some Properties of Georgian Propolis. Molecules, 27(22), 7714. 10.3390/molecules27227714

71. Ożarowski, M., Karpiński, T. M., Alam, R., & Łochyńska, M. (2022). Antifungal Properties of Chemically Defined Propolis from Various Geographical Regions. Microorganisms, 10(2), 364. 10.3390/microorganisms10020364

72. Pappas, P. G., Lionakis, M. S., Arendrup, M. C., Ostrosky-Zeichner, L., & Kullberg, B. J. (2018). Invasive candidiasis. Nature Reviews Disease Primers, 4, 18026. 10.1038/nrdp.2018.26

73. Paraje, M. G., Correa, S. G., Renna, M. S., Theumer, M., & Sotomayor, C. E. (2008). *Candida albicans*-secreted lipase induces injury and steatosis in immune and parenchymal cells. Canadian Journal of Microbiology, 54(8), 647–659. 10.1139/w08-048

74. Pasrija, R., Krishnamurthy, S., Prasad, T., Ernst, J. F., & Prasad, R. (2005). Squalene epoxidase encoded by *ERG1* affects morphogenesis and drug susceptibilities of *Candida albicans*. The Journal of Antimicrobial Chemotherapy, 55(6), 905–913. 10.1093/jac/dki112

75. Pfaffl M. W. (2001). A new mathematical model for relative quantification in real-time RT-PCR. Nucleic acids Research, 29(9), e45. 10.1093/nar/299.e45

76. Pippi, B., Lana, A. J., Moraes, R. C., Güez, C. M., Machado, M., de Oliveira, L. F., Lino von Poser, G., & Fuentefria, A. M. (2015). *In vitro* evaluation of the acquisition of resistance, antifungal activity and synergism of Brazilian red propolis with antifungal drugs on *Candida spp*. Journal of Applied Microbiology, 118(4), 839–850. 10.1111/jam.12746

77. Plaine, A., Walker, L., Da Costa, G., Mora-Montes, H. M., McKinnon, A., Gow, N. A., Gaillardin, C., Munro, C. A., & Richard, M. L. (2008). Functional analysis of *Candida albicans* GPI-anchored proteins: roles in cell wall integrity and caspofungin sensitivity. Fungal Genetics and Biology, 45(10), 1404–1414. 10.1016/j.fgb.2008.08.003

78. Qi, W., Acosta-Zaldivar, M., Flanagan, P. R., Liu, N. N., Jani, N., Fierro, J. F., Andrés, M. T., Moran, G. P., & Köhler, J. R. (2022). Stress- and metabolic responses of *Candida albicans* require Tor1 kinase N-terminal HEAT repeats. PLoS Pathogens, 18(6), e1010089. 10.1371/journal.ppat.1010089

79. R Core Team (2023). R: A language and environment for statistical computing. R Foundation for Statistical Computing, Vienna, Austria. https://www.R-project.org/

80. Reedy, J. L., Filler, S. G., & Heitman, J. (2010). Elucidating the *Candida albicans* calcineurin signaling cascade controlling stress response and virulence. Fungal Genetics and Biology, 47(2), 107–116. 10.1016/j.fgb.2009.09.002

81. Robbins, N., & Cowen, L. E. (2023). Roles of Hsp90 in *Candida albicans* morphogenesis and virulence. Current Opinion in Microbiology, 75, 102351. 10.1016/j.mib.2023.102351

82. Rodrigues M. L. (2018). The Multifunctional Fungal Ergosterol. mBio, 9(5), e01755–18. 10.1128/mBio.01755-18

83. Saavedra-Molina, A., Uribe, S., & Devlin, T. M. (1990). Control of mitochondrial matrix calcium: Studies using fluo-3 as a fluorescent calcium indicator. Biochemical and Biophysical Research Communications, 167(1), 148–153. 10.1016/0006-291x(90)91743-c

84. Sakai, Y., Murdanoto, A. P., Konishi, T., Iwamatsu, A., & Kato, N. (1997). Regulation of the formate dehydrogenase gene, FDH1, in the methylotrophic yeast *Candida boidinii* and growth characteristics of an FDH1-disrupted strain on methanol, methylamine, and choline. Journal of Bacteriology, 179(14), 4480–4485. 10.1128/jb.179.14.4480-4485.1997

85. Sanglard, D., McCallum, N., Gysler, C., Laloux, C., & Sanglard, C. (2021). Aequorin as a useful calcium-sensing reporter in Candida albicans. mSphere, 6(2), e01046–20. 10.1128/mSphere.01046-20

86. Schwartz, J. A., Olarte, K. T., Michalek, J. L., Jandu, G. S., Michel, S. L., & Bruno, V. M. (2013). Regulation of copper toxicity by *Candida albicans* GPA2. Eukaryotic Cell, 12(7), 954–961. 10.1128/EC.00344-12

87. Shahina, Z., Ndlovu, E., Persaud, O., Sultana, T., & Dahms, T. E. S. (2022). *Candida albicans* Reactive Oxygen Species (ROS)-Dependent Lethality and ROS-Independent Hyphal and Biofilm Inhibition by Eugenol and Citral. Microbiology Spectrum, 10(6). 10.1128/spectrum.03183-22

88. Silva-Beltrán, N. P., Boon, S. A., Ijaz, M. K., McKinney, J., & Gerba, C. P. (2023). Antifungal activity and mechanism of action of natural product derivates as potential environmental disinfectants. Journal of Industrial Microbiology & Biotechnology, 50(1). 10.1093/jimb/kuad036

89. Silva-Carvalho, R., Baltazar, F., & Almeida-Aguiar, C. (2015). Propolis: A Complex Natural Product with a Plethora of Biological Activities That Can Be Explored for Drug Development. Evidence-Based Complementary and Alternative Medicine, 2015, 206439. 10.1155/2015/206439

90. Slowikowski, K. (2023). ggrepel: Automatically Position Non-Overlapping Text Labels with ‘ggplot2’. R package version 0.9.4. https://CRAN.R-project.org/package=ggrepel

91. Sudbery, P., Gow, N., & Berman, J. (2004). The distinct morphogenic states of *Candida albicans*. Trends in Microbiology, 12(7), 317–324. 10.1016/j.tim.2004.05.008

92. Sun, C., Zhang, M., Niu, C., Wang, C., Wang, Y., & Chen, L. (2022). A Cecropin-4 derived peptide C18 inhibits *Candida albicans* by disturbing intracellular Ca²⁺ homeostasis and cell integrity. Frontiers in Microbiology, 13, 872322. 10.3389/fmicb.2022.872322

93. Sun, Z., Kuczek, T., & Zhu, Y. (2014). Statistical calibration of qRT-PCR, microarray and RNA-Seq gene expression data with measurement error models. The Annals of Applied Statistics, 8(2). 10.1214/14-aoas721

94. Swenson, K. A., Min, K., & Konopka, J. B. (2024). *Candida albicans* pathways that protect against organic peroxides and lipid peroxidation. PLoS Genetics, 20(10), e1011455. 10.1371/journal.pgen.1011455

95. Talapko, J., Juzbašić, M., Matijević, T., Pustijanac, E., Bekić, S., Kotris, I., & Škrlec, I. (2021). *Candida albicans* - The Virulence Factors and Clinical Manifestations of Infection. Journal of Fungi, 7(2), 79. 10.3390/jof7020079

96. Tobaldini-Valerio, F. K., Bonfim-Mendonça, P. S., Rosseto, H. C., Bruschi, M. L., Henriques, M., Negri, M., Silva, S., & Svidzinski, T. I.e (2016). Propolis: a potential natural product to fight *Candida* species infections. Future Microbiology, 11, 1035–1046. 10.2217/fmb-2015-0016

97. UniProt Consortium (2025). UniProt: the Universal Protein Knowledgebase in 2025. Nucleic acids research, 53(D1), D609–D617. 10.1093/nar/gkae1010

98. Wang, H., Liang, Y., Zhang, B., Zheng, W., Xing, L., & Li, M. (2011). Alkaline stress triggers an immediate calcium fluctuation in *Candida albicans* mediated by Rim101p and Crz1p transcription factors. FEMS Yeast Research, 11(5), 430–439. 10.1111/j.1567-1364.2011.00730.x

99. Warnes, G.R., Bolker, B., Bonebakker, L., Gentleman, R., Huber, W., Liaw, A., et al. (2022). gplots: Various R Programming Tools for Plotting Data. R package version 3.1.3. https://CRAN.R-project.org/package=gplots

100. Watanabe, H., Azuma, M., Igarashi, K., & Ooshima, H. (2005). Analysis of Chitin at the Hyphal Tip of *Candida albicans* Using Calcofluor White. Bioscience, Biotechnology, and Biochemistry, 69(9), 1798–1801. 10.1271/bbb.69.1798

101. Whaley, S. G., Tsao, S., Weber, S., Zhang, Q., Barker, K. S., Raymond, M., & Rogers, P. D. (2016). The RTA3 Gene, Encoding a Putative Lipid Translocase, Influences the Susceptibility of *Candida albicans* to Fluconazole. Antimicrobial Agents and Chemotherapy, 60(10), 6060–6066. 10.1128/AAC.00732-16

102. Whaley, S. G., Zhang, Q., Caudle, K. E., & Rogers, P. D. (2018). Relative Contribution of the ABC Transporters Cdr1, Pdh1, and Snq2 to Azole Resistance in Candida glabrata. Antimicrobial Agents and Chemotherapy, 62(10), e01070–18. 10.1128/AAC.01070-18

103. WHO. (2022). Fungal Priority Pathogens List to Guide Research, Development and Public Health Action. World Health Organization, Geneva. Licence: CC BY-NC-SA 3.0 IGO.

104. Wickham, H. (2016). ggplot2: Elegant Graphics for Data Analysis. Springer International Publishing. 10.1007/978-3-319-24277-4

105. Wilke, C.O. (2020). cowplot: Streamlined Plot Theme and Plot Annotations for ‘ggplot2’. R package version 1.2.0. https://wilkelab.org/cowplot/

106. Williams, R. B., & Lorenz, M. C. (2020). Multiple Alternative Carbon Pathways Combine To Promote *Candida albicans* Stress Resistance, Immune Interactions, and Virulence. mBio, 11(1), e03070–19. 10.1128/mBio.03070-19

107. Wu, T., Hu, E., Xu, S., Chen, M., Guo, P., Dai, Z., Feng, T., Zhou, L., Tang, W., Zhan, L., Fu, X., Liu, S., Bo, X., & Yu, G. (2021). clusterProfiler 4.0: A universal enrichment tool for interpreting omics data. The Innovation, 2(3), 100141. 10.1016/j.xinn.2021.100141

108. Xiao, J., & Kai, G. (2012). A review of dietary polyphenol-plasma protein interactions: characterization, influence on the bioactivity, and structure-affinity relationship. Critical Reviews in Food Science and Nutrition, 52(1), 85–101. 10.1080/10408398.2010.499017

109. Xie, F., Wang, J., & Zhang, B. (2023). RefFinder: a web-based tool for comprehensively analyzing and identifying reference genes. Functional & Integrative Genomics, 23(2), 125. 10.1007/s10142-023-01055-7

110. Yardımcı, B. K., Şimşek Özek, N., Güler, A., Akköylü, A., & Çokak, T. A. (2025). Propolis impairs cellular proliferation by promoting oxidative damage and disrupting macromolecular composition in a yeast model. Scientific Reports, 15(1), 43085. 10.1038/s41598-025-27190-8

111. Ye, J., Coulouris, G., Zaretskaya, I., Cutcutache, I., Rozen, S., & Madden, T. L. (2012). Primer-BLAST: a tool to design target-specific primers for polymerase chain reaction. BMC Bioinformatics, 13, 134. 10.1186/1471-2105-13-134

112. Yu, G. (2023). enrichplot: Visualization of Functional Enrichment Result. R package version 1.22.0. https://bioconductor.org/packages/enrichplot

113. Zhang, P., Li, H., Cheng, J., Sun, A. Y., Wang, L., Mirchevska, G., Calderone, R., & Li, D. (2018). Respiratory stress in mitochondrial electron transport chain complex mutants of *Candida albicans* activates Snf1 kinase response. Fungal Genetics and Biology, 111, 73–84. 10.1016/j.fgb.2017.11.002

114. Zhao, X., Oh, S.-H., & Hoyer, L. L. (2007). Deletion of *ALS5*, *ALS6* or *ALS7* increases adhesion of *Candida albicans* to human vascular endothelial and buccal epithelial cells. Medical Mycology, 45(5), 429–434. 10.1080/13693780701377162

115. Ziegler, L., Terzulli, A., Gaur, R., McCarthy, R., & Kosman, D. J. (2011). Functional characterization of the ferroxidase, permease high-affinity iron transport complex from *Candida albicans*. Molecular Microbiology, 81(2), 473–485. 10.1111/j.1365-2958.2011.07704.x

