## Supplement for "Deciphering the antifungal mechanism of Polish ethanolic extracts of propolis against *Candida albicans*: Evidence for a multi-target mode of action"

**Supplementary materials**

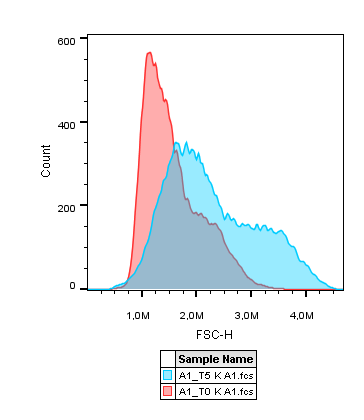

Figure S1. FSC-H measurement of cells grown overnight in YPD medium (non-inducing conditions) and cells incubated for 3 h in RPMI medium (hyphae-inducing conditions). An empirical FSC-H threshold was established based on comparison of signal distributions to distinguish blastospore population (signals below AU = 1.9) from the population enriched in hyphal/pseudohyphal forms (signals above AU = 1.9).

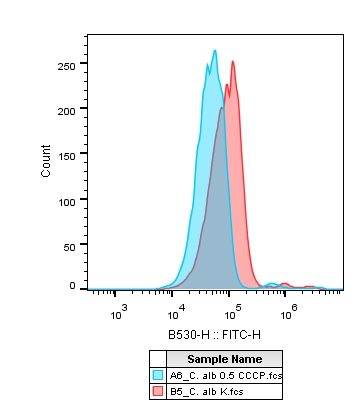

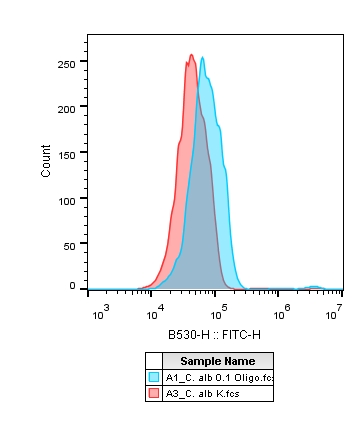

Figure S2. Fluorescence shift of CCCP- (left) and oligomycin A-treated *C. albicans* cells (right) in relation to untreated cells.

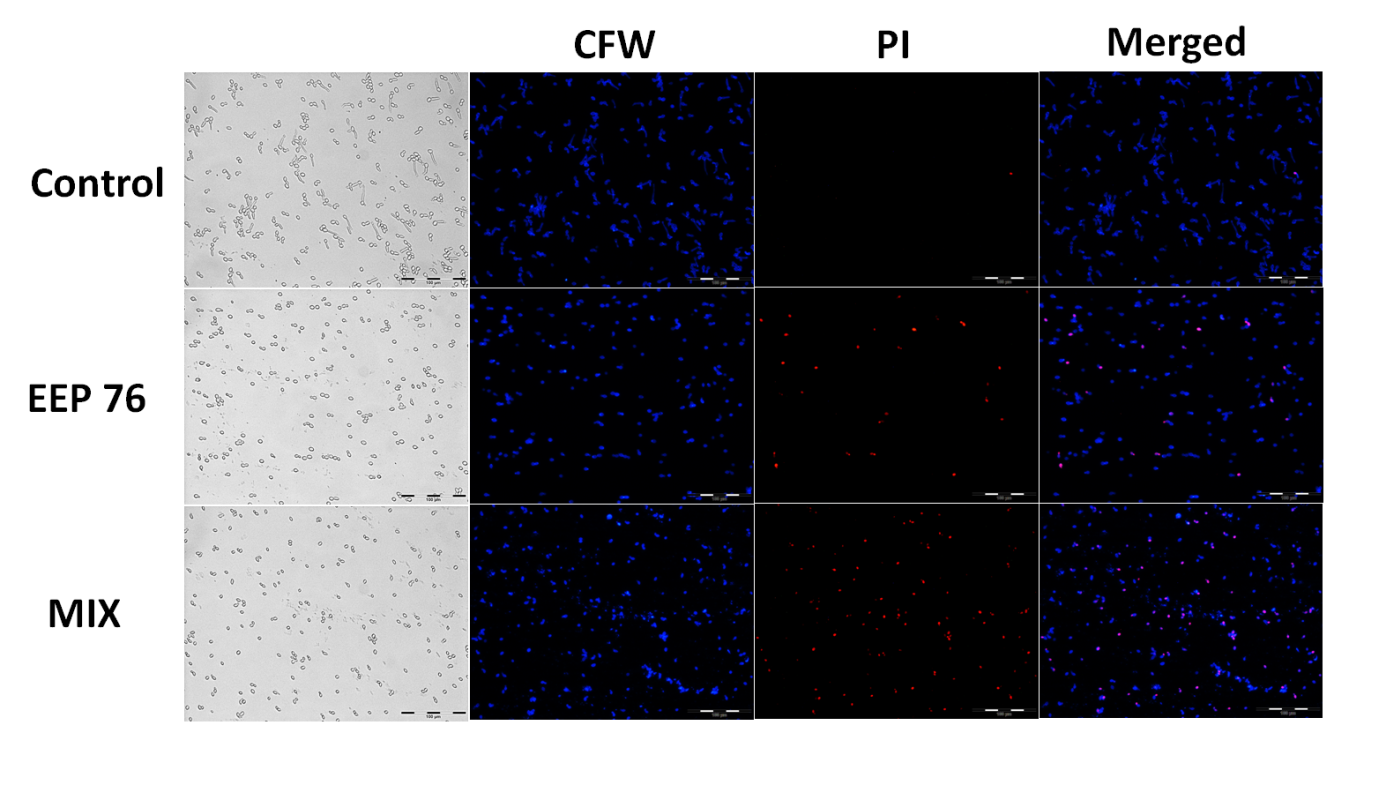

Figure S3. Bright-field and fluorescent microscopy pictures (20x magnification) of *C. albicans* cells treated with EtOH/DMSO (control), EEP 76, and MIX co-stained using Calcofluor White (CFW) and iodine propide (PI) dyes.

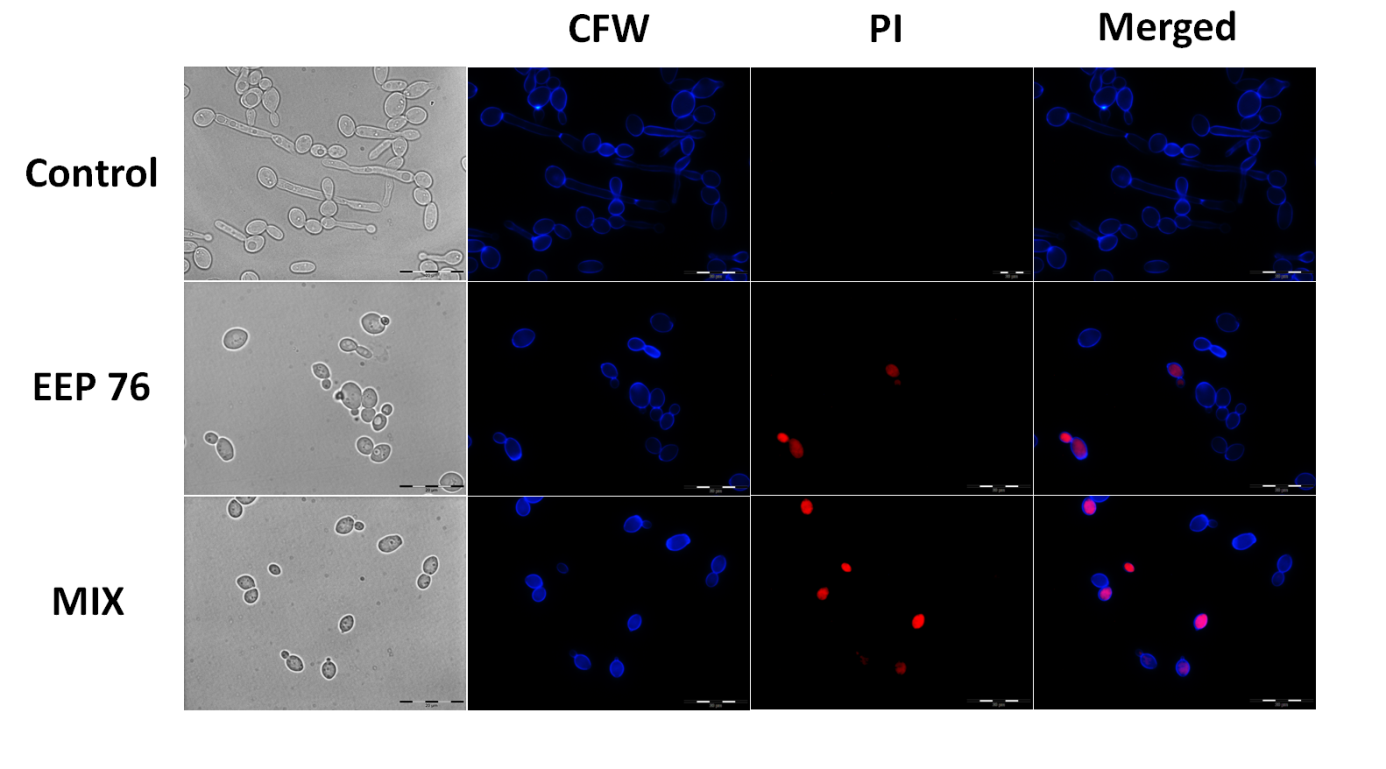

Figure S4. Bright-field and fluorescent microscopy pictures (100x magnification) of *C. albicans* cells treated with EtOH/DMSO (control), EEP 76, and MIX co-stained using Calcofluor White (CFW) and

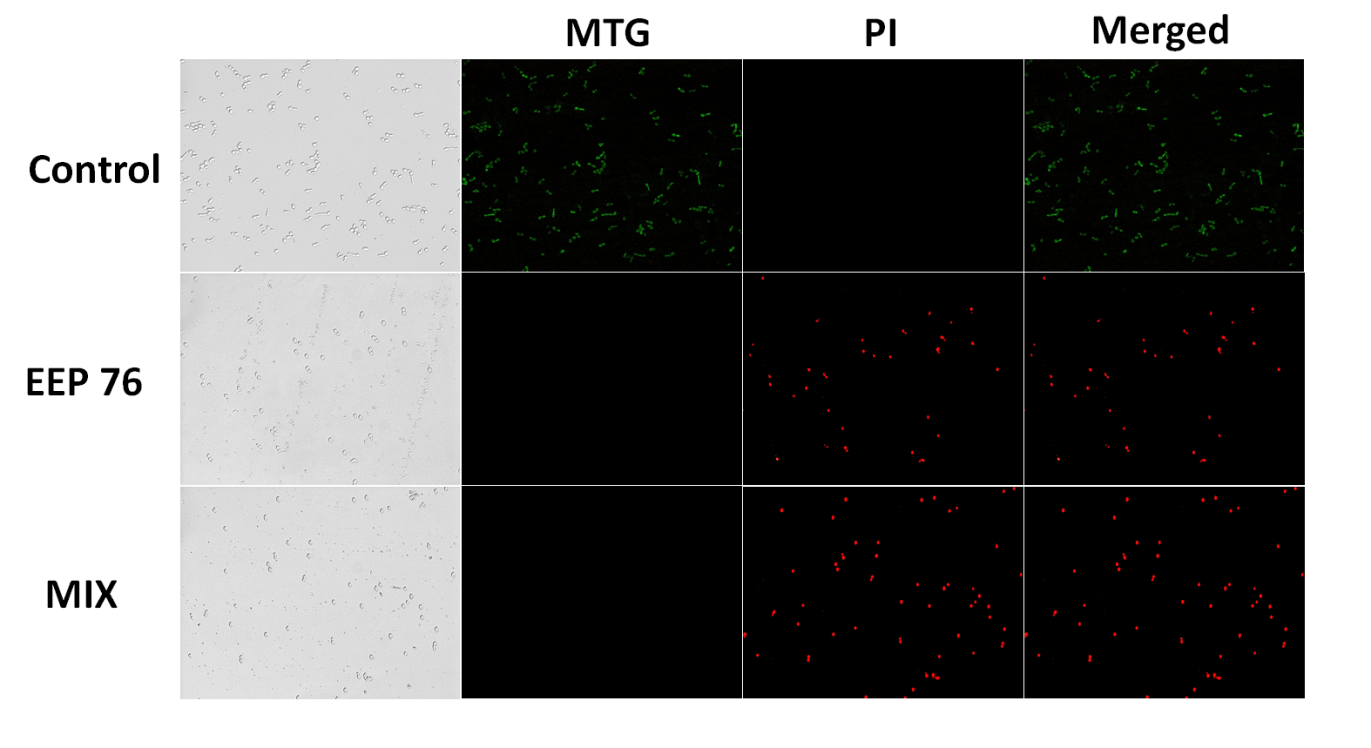

Figure S5. Bright-field and fluorescent microscopy pictures (20x magnification) of *C. albicans* cells treated with EtOH/DMSO (control), EEP 76, and MIX co-stained using Mitotracker™ Green (MTG) and iodine propide (PI) dyes.

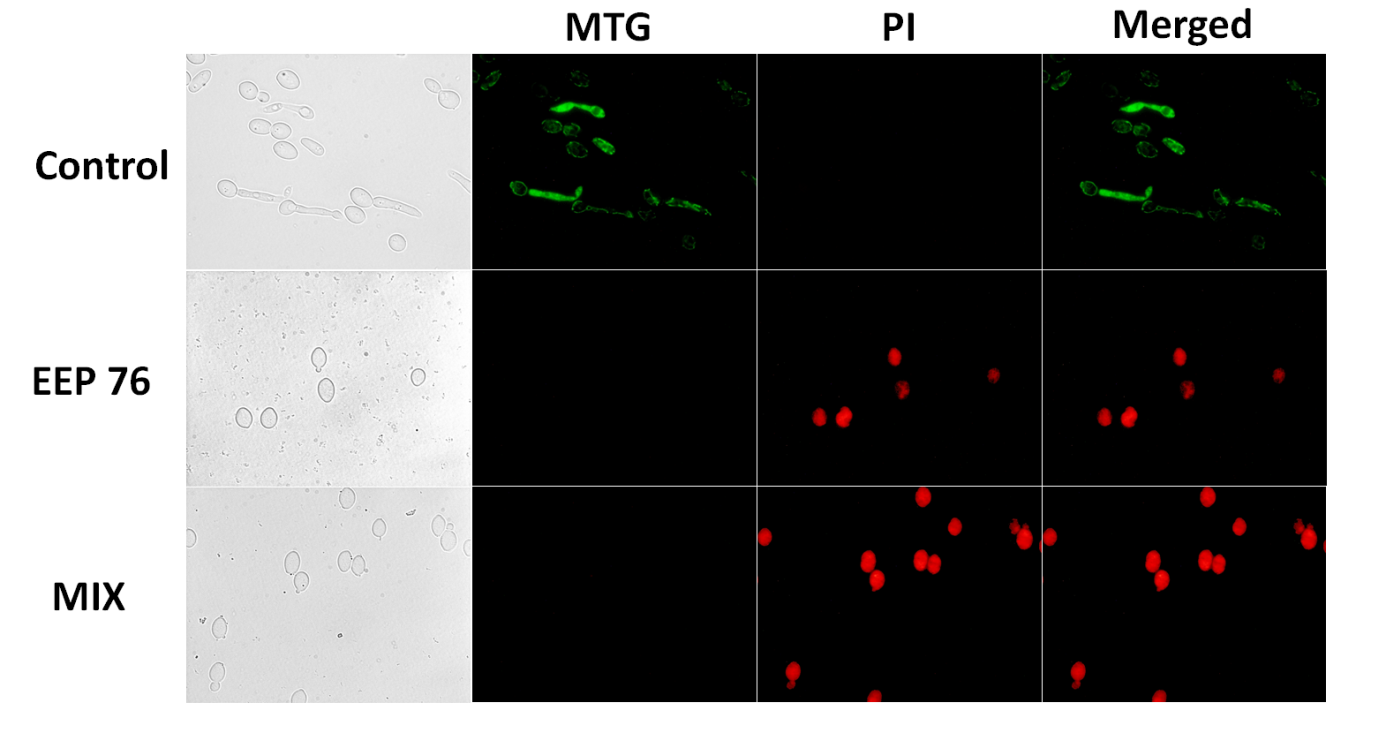

Figure S6. Bright-field and fluorescent microscopy pictures (100x magnification) of *C. albicans* cells treated with EtOH/DMSO (control), EEP 76, and MIX co-stained using Mitotracker™ Green (MTG) and iodine propide (PI) dyes.

Table S1. qPCR primer sequences and experimentally optimized concentrations used in experiments.

| **Gene** | **Primer sequence** | | **Optimized concentration [µM]** |
| --- | --- | --- | --- |
|  | **Forward (5' - 3')** | **Reverse (5' - 3')** |  |
| ERG1 | TTCCTGATGCCGTTGATGGT | AAATGCAGCCCCACGTACTC | 0.2 |
| LIP3 | CCGAACCATTCTGGGGAACA | CCATCCAACGAGCCGTGATA | 0.3 |
| SAP4 | CCGTTGGTATTGGTGGTGTTTC | TGGTGGCTTCGTTGCTTTG | 0.45 |
| CDR1 | TGCTGCCATGTTCTTTGCTG | CGACAATTGGTCTGGCTTCG | 0.05 |
| CDR2 | GCAATTGGTGATGGGCCTTA | AAAGTGCTCCACCTCGGAAA | 0.2 |
| HSP12 | TGCAGACAACGTGCAACAAG | TGAGCTGTTTCAGCAAGGGT | 0.3 |
| FLU1 | TGCTTGGTTCACGAGATCCTT | AAGGCAGCAAGACCAACAGA | 0.3 |
| COB | TTAGGGTAGGTCGGGTGGAT | GTGGTGCAGTCTGTAGCGATT | 0.45 |
| SNQ2 | TCGGGGTTGAATTGGTTGCT | TGGCCCAAGCAGATTGTGAA | 0.2 |
| NAD1 | TATGCAACGGAGAGTTGGTCC | AGCTGCATCAGCAATAGCCA | 0.1 |
| STP1 | CAGTCGGAGGGACACTTTCA | AACAGCCAGCACATCTTCCA | 0.3 |
| RBR1 | TGGTGCTTCATCTGCTACCG | AATGGCTGCCAAGCTACCAG | 0.3 |
| ALS1 | TTCTCATGAATCAGCATCCACAA | CAGAATTTTCACCCATACTTGGTTTC | 0.4 |
| ALS3 | CAACTTGGGTTATTGAAACAAAAACA | AGAAACAGAAACCCAAGAACAACCT | 0.2 |
| ALS7 | GAAGAGAACTAGCGTTTGGTCTAGTTGT | GCGACATGGAAAGTCTTTGACTAAC | 0.4 |
| HWP1 | GACCGTCTACCTGTGGGACAGT | GCTCAACTTATTGCTATCGCTTATTACA | 0.2 |
| ACT1 | TGATGATGTTGCTAGATTATGGTCG | TCGTCACGGGTATTTTCTCTTGT | 0.2 |
| RIP1 | AACCACCACCACCTTATCCAA | AGTTTCTGCTTCCTTGACCG | 0.3 |
| PMA1 | TTGCTTATGATAATGCTCCATACGA | TACCCCACAATCTTGGCAAGT | 0.05 |
| LSC2 | CGTCAACATCTTTGGTGGTATTGT | TTGGTGGCAGCAATTAAACCT | 0.3 |
| RPP2B | GGTGTTGAAGCCGAAGAATCC | TGGTGTTACCTTCAGCAATCAA | 0.1 |

Table S2. Expression stability ranking of the reference genes according to RefFinder software for samples exposed to EEP.

| **Method** | **3 h exposure to EEP** | | | | |
| --- | --- | --- | --- | --- | --- |
|  | **Ranking order^a^** | | | | |
|  | **1** | **2** | **3** | **4** | **5** |
| BestKeeper | *act1* | *rip1* | *lsc2* | *rpp2b* | *pma1* |
| Normfinder | *lsc2* | *act1* | *rip1* | *rpp2b* | *pma1* |
| Genorm | *rpp2b* \| *rip1* | *-* | *act1* | *lsc2* | *pma1* |
| delta-Ct | *act1* | *lsc2* | *rip1* | *rpp2b* | *pma1* |
| **Recommended comprehensive ranking** | *act1* | *rip1* | *lsc2* | *rpp2b* | *pma1* |
| ^a^ 1, the best among the studied genes; 5, the worst among the studied genes. | | | |  |  |

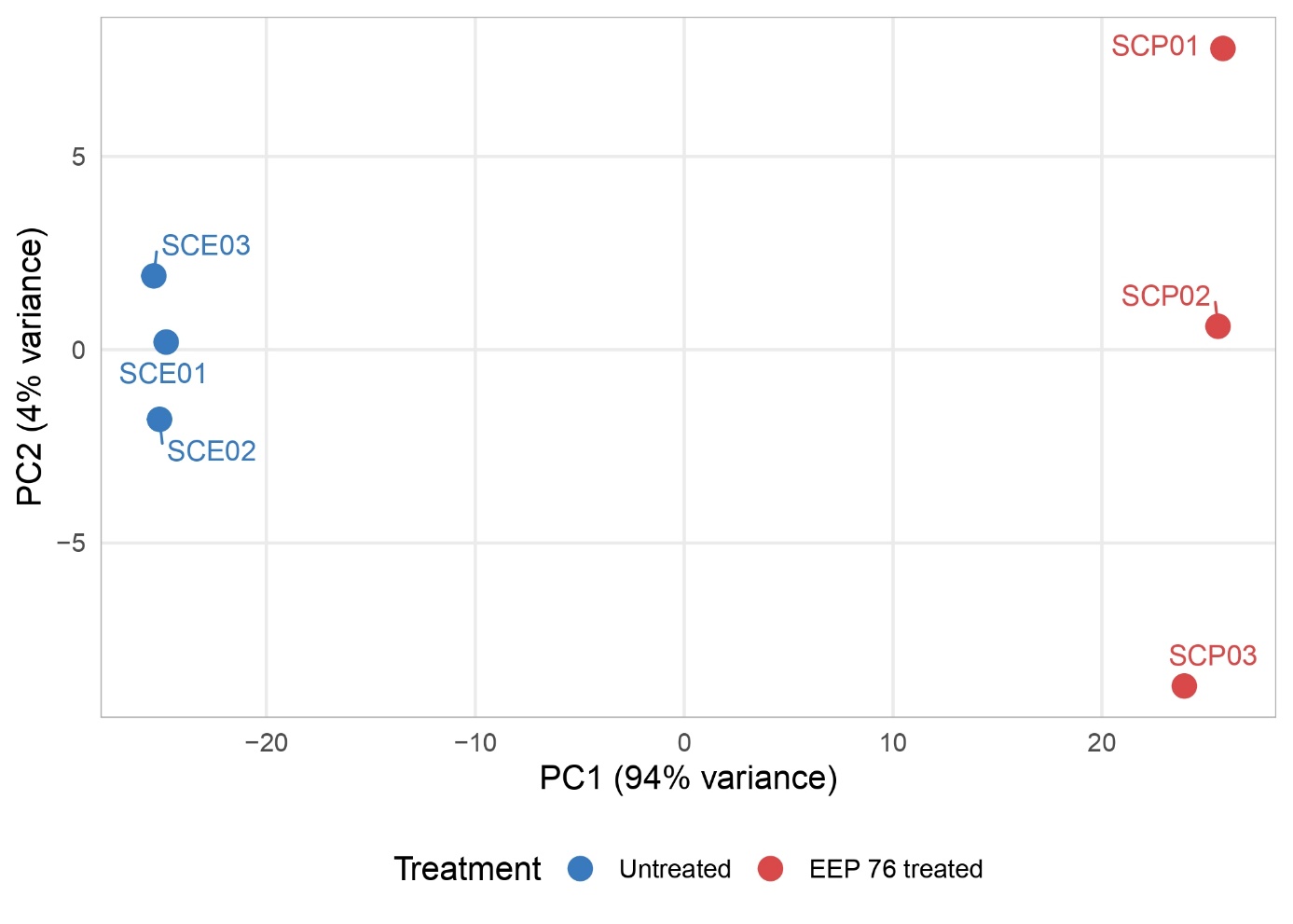

Figure S7. Principal component analysis (PCA) of RNA-seq samples. PCA was performed on regularized log-transformed (rlog) expression values from DESeq2. Each point represents one biological replicate: untreated control (SCE01–SCE03) and EEP 76 treated (SCP01–SCP03). Axes show the first two principal components with the percentage of variance explained.

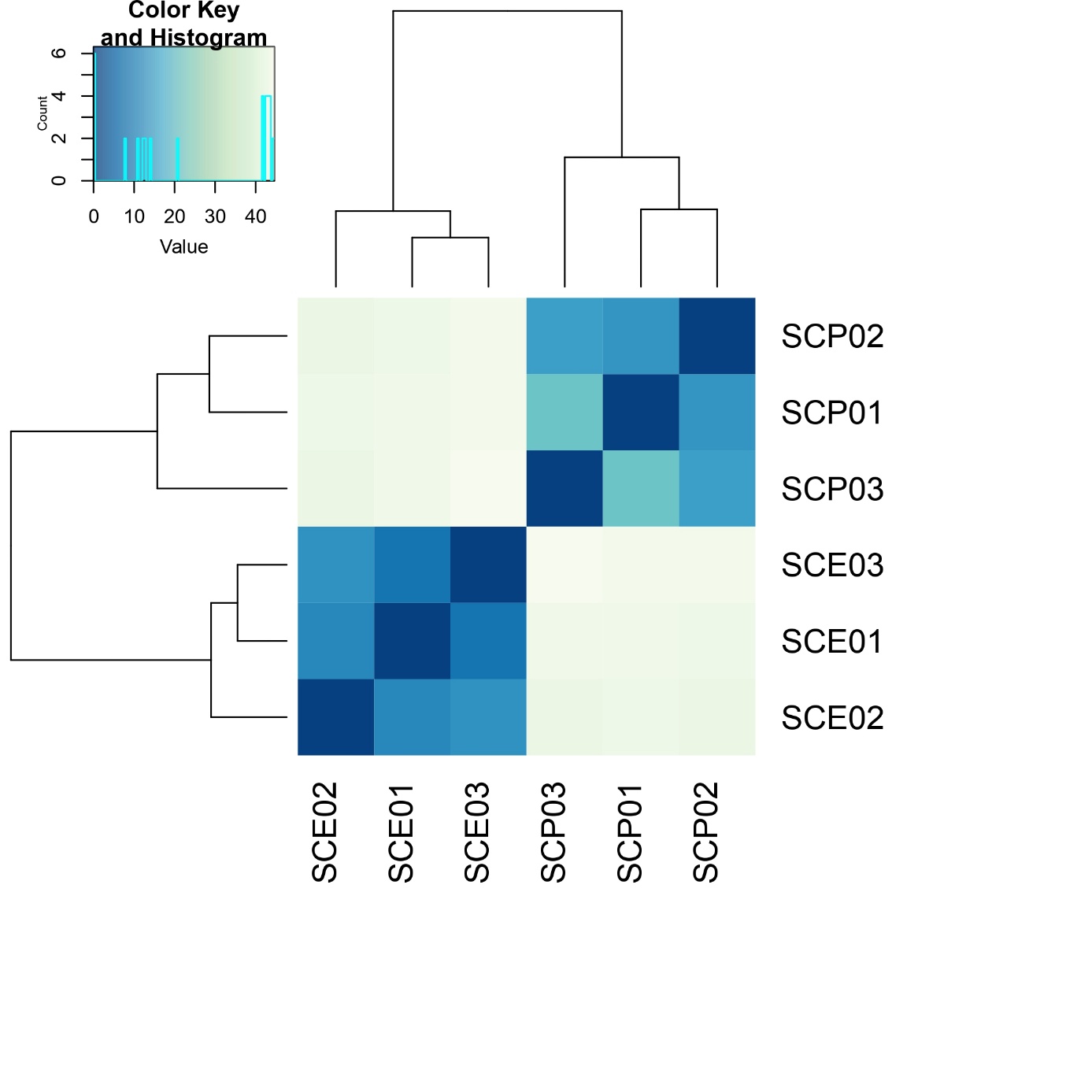

Figure S8. Sample-to-sample distance heatmap. Euclidean distances were computed on rlog-transformed expression values and visualized with hierarchical clustering. Color intensity reflects pairwise distance between samples, with darker shades indicating greater similarity.
